# Comprehensive Mapping of Immune Nanobody Repertoires with NanoMAP

**DOI:** 10.64898/2026.03.05.709882

**Authors:** William L. White, Edward Moseley, Jacqueline M. Tremblay, Jackson Reilly, Akram A. Da’dara, Patrick J. Skelly, Lenore J. Cowen, Charles B. Shoemaker

## Abstract

Nanobodies have recently emerged as alternatives to classical antibodies in therapeutic and diagnostic contexts from parasites to bacteria to viruses, promising improved stability and simpler manufacturing. To improve nanobody discovery efficiency, we developed an integrated experimental and computational pipeline for detailed characterization of the target binding properties of complete alpaca immune repertoires using our custom Nanobody Meta-clustering Analysis Platform (NanoMAP). We tested our pipeline on three distinct pools of targets, immunizing two alpacas with each pool and generating cDNA and phage display libraries from their immune repertoires. We then panned the phage libraries on each target. To produce more detailed binding information, we performed panning variations using subunits, natural variants, intact pathogens, and binding site competitors. Deep sequencing reads from nanobody libraries before and after each panning were pooled and analyzed with NanoMAP to identify nanobody clonal families and assess their levels of enrichment from the library in each panning, reflecting their affinities. NanoMAP outperformed standard clustering methods, producing clonal families that are coherent in sequence and function and detecting rare but high affinity families. By aggregating sequencing data within clonal families, NanoMAP produced reliable and rich data on nanobody repertoire binding phenotypes for each antigen, enhancing nanobody discovery capabilities.

## 1 Introduction

The high affinity and specificity of conventional antibodies has made their development into therapeutics and diagnostics highly successful in a wide range of human diseases [14, 20, 28]. Nanobodies (V_H_ domains of heavy-chain-only antibodies, V_H_Hs) offer the same potential for binding affinity and specificity as conventional antibodies, with significant improvements in manufacturing, storage, and stability [1]. Nanobodies also allow more flexibility in their design as they can be linked to each other or to other fusion partners, improving binding properties such as affinity and polyspecificity, and enhancing other applications such as diagnostic testing [2, 22]. These improved properties have led to rapid growth in literature describing the development of nanobodies into therapeutics and diagnostics as a productive endeavor [10, 11, 39]. The success of nanobody therapeutics and diagnostics highlights their relevance and utility while underscoring the need to develop improved methods for nanobody library selection and computational analysis.

Although many groups have developed and refined methods for immunization, screening, and analysis of nanobody repertoires, a generalizable pipeline does not exist to comprehensively characterize nanobody immune repertoires targeting arbitrary antigens of interest. Most prior work has identified candidates in low-throughput by randomly selecting a small sampling of nanobodies from a target-enriched phage display library, with a few recent exceptions that used high-throughput sequencing (HTS) to evaluate enrichment across the entire library [21, 34, 36]. But even these HTS experiments focus on a single target at a time, and provide limited binding information for each nanobody in the library. Additionally, previous analysis methods focus on individual nanobody sequences, thereby missing out on trends within clonal families of nanobodies derived from the same initial B-cell clone. To address these issues, we developed an integrated experimental and computational pipeline (**fig. 1**) that takes a set of antigens of interest as input, and outputs binding strength, epitope, and variant specificity information for nearly all nanobody clonal families in the alpaca immune repertoire. Critically, our pipeline relies on sequence-based clustering to aggregate data within clonal families. We tested two well-established clustering methods, SCOPer [25] (from the Immcantation package) and MMseqs2 [30], against our own clustering methods and identified optimal clustering parameters for each method. Our Nanobody Meta-clustering Analysis Platform (NanoMAP) includes our clustering methods, but can be used with any clustering method to analyze sequencing data generated from immunization and panning experiments. Using NanoMAP, we are now able to provide far more information about many more nanobodies, and enable efficient selection of candidate therapeutics and diagnostics with optimal, purpose-suited, properties based on a variety of relevant characteristics.

**Figure 1:**
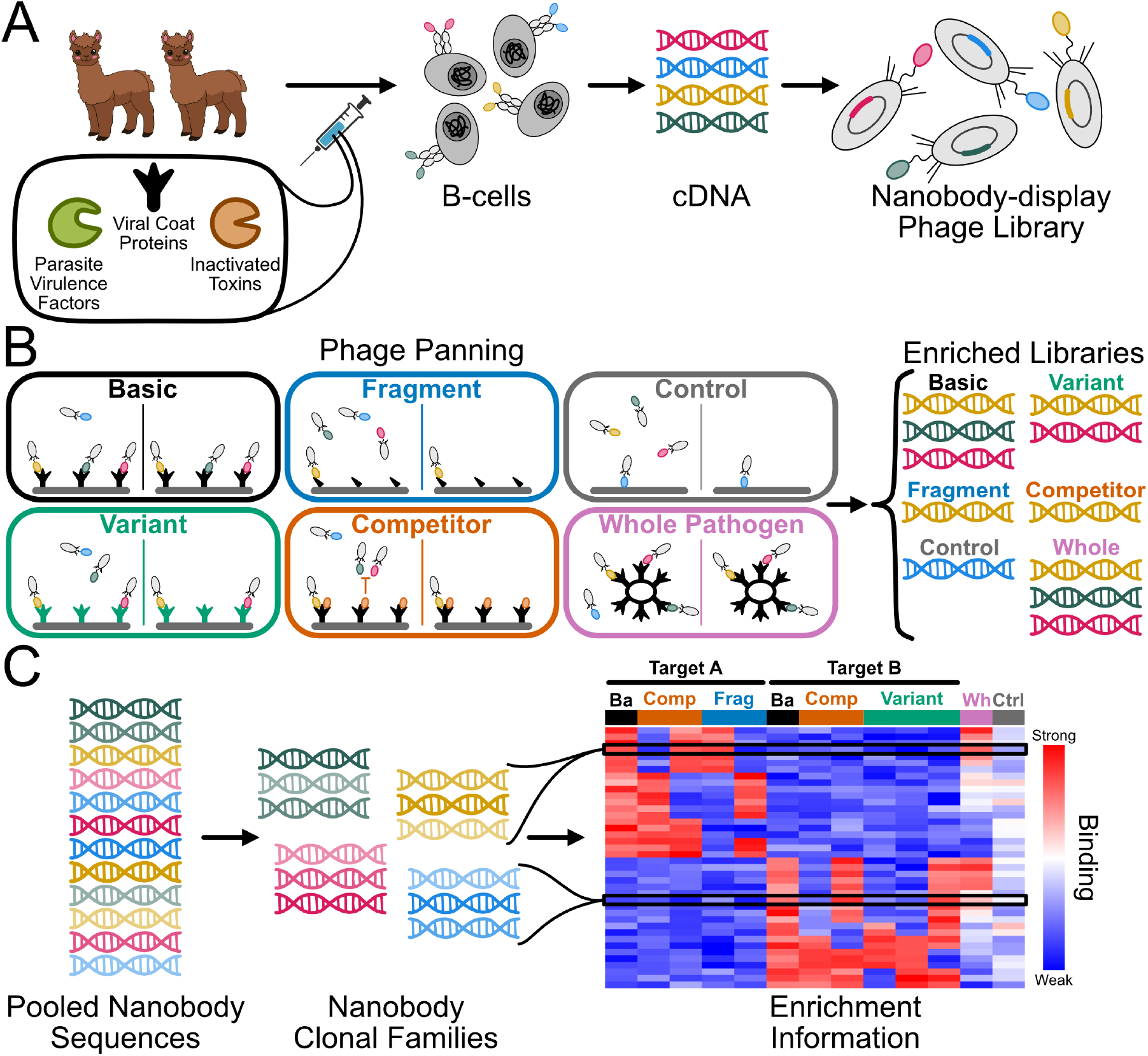
Alpaca immunization and nanobody phage display facilitate discovery of nanobodies binding diverse targets. Schematics showing an outline of the experimental procedures. A) Alpacas are immunized over several weeks, using a pool of multiple distinct target proteins. After immunization, a blood sample is collected and nanobody cDNA is extracted and converted into a phage display library. B) Phage display panning experiments are performed in several possible variations using the phage library. In a basic panning, phage are washed over a target-coated surface and unbound phage are washed off, leaving the bound phage to be collected and sequenced. Control panning experiments are performed in the same way as basic panning, but in the absence of the target. Fragment, variant, competitor, and whole pathogen panning experiments are performed identically to the basic panning, either using a fragment or variant of the target, blocking binding to a specific site using a competitor, or using the whole pathogen to capture phage, respectively. C) The initial display library and each panned library are barcoded, and pooled for sequencing. Sequencing data from all panning experiments are pooled and clustered into clonal families. Read count data are aggregated within clonal families and compared across panning experiments to determine the binding profile of each family.

Our pipeline consists of three major steps, and offers both experimental and computational improvements over existing pipelines. First, alpacas are immunized with a pool of antigens (**fig. 1A**), facilitating parallel nanobody discovery campaigns. The resulting immune repertoires are then converted to phage display libraries using established methods [16, 27]. Second, phage libraries are panned for binding under a variety of conditions, including in the presence of binding site competitors, or using naturally occurring variants of the target (**fig. 1B**). Third, the resulting libraries are sequenced and clustered into clonal families. In combination, the panning conditions and clustering provide rich information about the binding phenotypes present in the repertoire that are often overlooked when only standard panning is performed. Although others have reported similar approaches [21, 34, 36], they have not been combined in a manner that allows the rich and comprehensive characterization reported here. NanoMAP (**fig. 1C, fig. 2**) substantially improves the detection of rare binding phenotypes by aggregating data across multiple related nanobodies and provides a more succinct representation of the immune repertoire, improving candidate selection.

**Figure 2:**
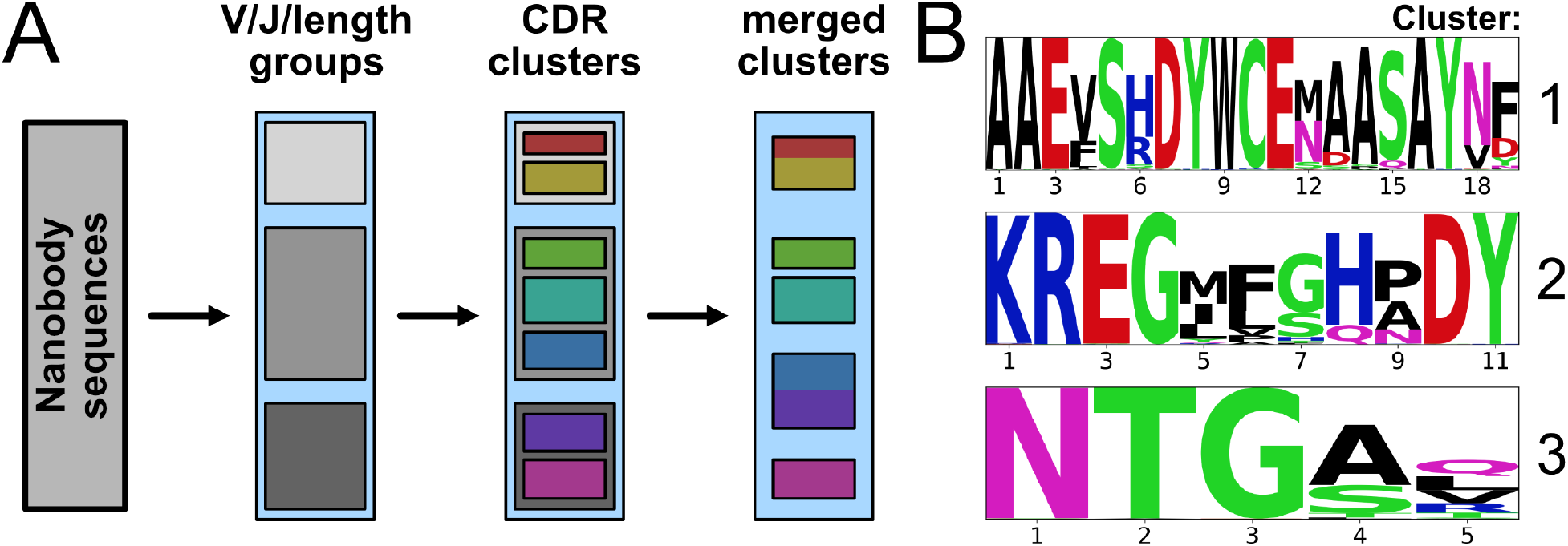
NanoMAP clustering allows flexible clustering through a merging step. A) Schematic of the NanoMAP clustering method. Sequences are first broken into groups that share V and J segment annotations and CDR lengths, then each group is refined into subgroups of similar sequences by hierarchical clustering based on CDR sequences. The fine-grained groups are then compared to each other and merged if they are sufficiently similar, allowing for minor differences in CDR lengths. The resulting clusters are referred to as clonal families. B) Logo plots of the CDR3 sequences from three representative clonal families.

## 2 Results

### 2.1 NanoMAP clustering yields highly accurate clonal families

To equitably compare NanoMAP to currently accepted clustering methods (Immcantation’s SCOPer [25] and MMseqs2 [30], we tested a range of parameter settings and identified optimal parameters for each method (**section 4.6**). Each clustering method showed large variations across all scores as the linkage method and distance cutoffs were varied (**fig. S1-11**), underscoring the importance of choosing the optimal parameters. We decide optimality based on four different scores we compute on the clustering: one of the four scores is the standard silhouette score, one requires gold standard hand-curated clusterings, but two are new measures of cluster quality that we think are of independent interest (**section 4.6**).

To evaluate the overall performance of each clustering method, we used a leave-one-out (LOO) cross-validation approach (**section 4.6.6, fig. 3**). We found that MMseqs2, although it is significantly faster and requires less memory (**table S4**), achieves much worse Silhouette, phenotypic quality, and ARI scores, likely due to the fact that it was not created for nanobody clustering specifically. NanoMAP and SCOPer methods show more similar performance, with NanoMAP outperforming SCOPer more often than not. Interestingly, despite the known benefits of aggregate methods like meta-clustering [32], NanoMAP individual clusterings (**section S1.2.1**) outperformed meta-clusterings (**section S1.2.2**) in nearly all datasets and metrics on our selected test datasets. Even in the stability score, which was expected to improve in meta-clustering through the use of many individual clustering parameter settings, NanoMAP individual clustering was comparable to or better than the meta-clustering, suggesting that optimal clustering parameters do not vary significantly with dataset size. Additionally, NanoMAP individual and SCOPer clusterings each produced the same optimal parameters in all folds of the LOO analysis, while MMseqs2 and NanoMAP meta-clustering each produced two distinct optimal parameter settings, depending on the datasets used. We discuss how these results impact our recommended parameter settings for NanoMAP in **section 3**.

**Figure 3:**
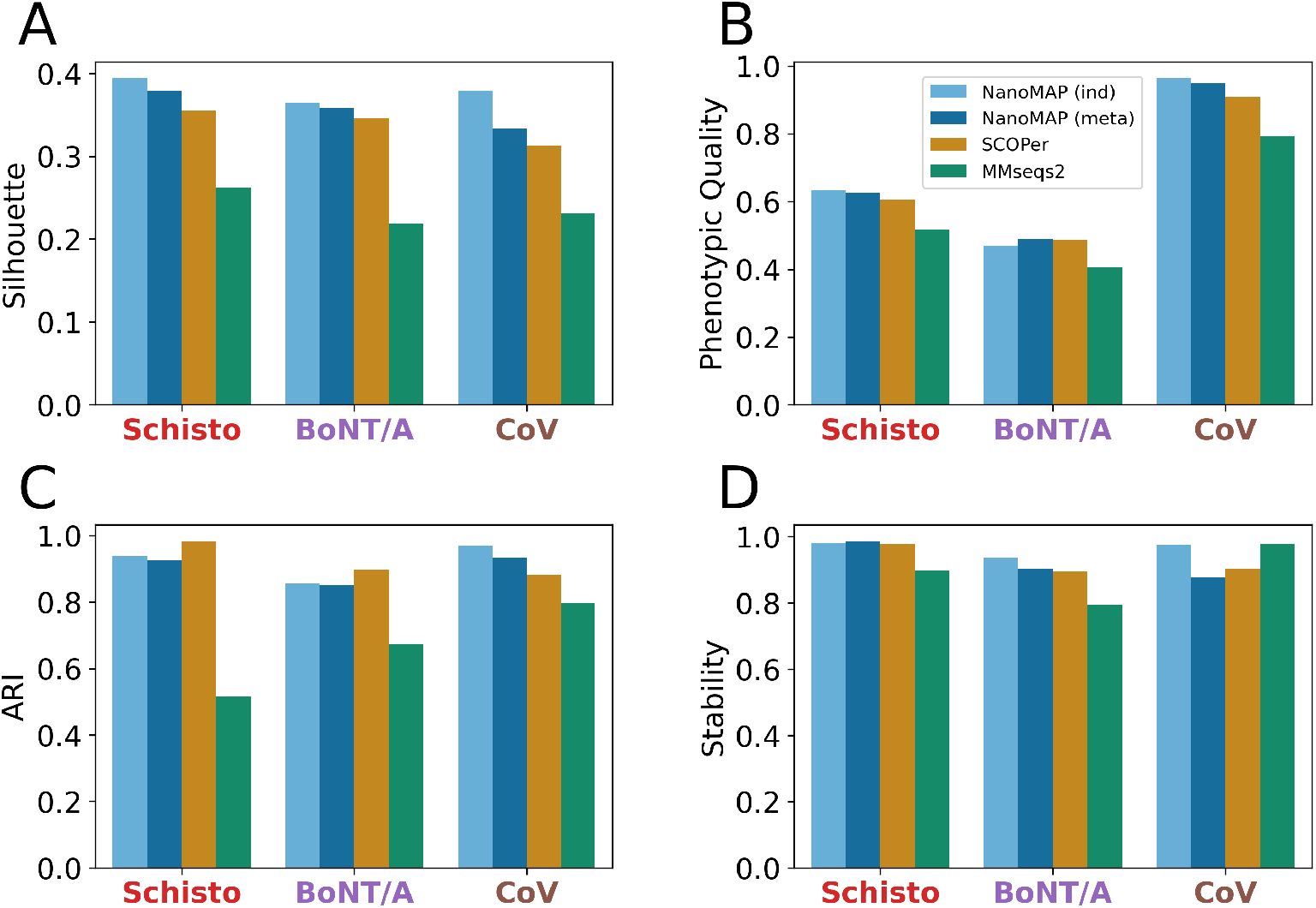
NanoMAP produces high-quality and consistent clusterings. Bar charts showing the performance of NanoMAP and existing clustering methods across all three datasets, using optimal parameter settings. A) Non-normalized silhouette scores calculated using CDR distance. B) Phenotypic quality scores, measuring the consistency of within-cluster binding phenotypes. C) ARI scores comparing each clustering to an expert-curated clustering of a subset of the sequences. D) Down-sampling stability scores, reporting the ARI of the down-sampled clustering that is least similar to the fill-size clustering.

### 2.2 Validation of Immune Repertoire Sampling and Panning

Before analyzing the behavior of each clonal family in each panning experiment, we applied several techniques, described below, to evaluate the quality of the data produced in these experiments.

#### 2.2.1 Library preparation methods produce minimal sequence errors

First, we evaluated the error rate of our DNA preparation and sequencing methods to ensure that the data we collected accurately represented the samples they were derived from. We prepared a single clonal plasmid as if it were a DNA library and sequenced the resulting sample using the same methods as for the phage display samples. Because the composition of the sample was known, we could determine the location and number of mutations that existed in each read. We found that the majority (76%) of reads had no mutations, with very few (*<*7%) containing more than 10 mutations (**fig. S13A**), resulting in an overall error rate of 0.34 errors per read. We also found a subset (representing roughly 19% of all reads) of error-containing sequences that were observed many times (**fig. S13B**), indicating that they likely arose during the sample preparation, rather than from sequencing noise. We additionally examined the distribution of positions that were mutated, and found relatively even distribution throughout the length of the sequence, with elevated rates of mutation only at the first and last positions (**fig. S13C**). Given the relatively low rate of and even distribution of mutations, it is likely that mutated sequences will be clustered in the same clonal family as the sequence they were derived from, minimally impacting the interpretation of our experimental results.

#### 2.2.2 Sequencing results are reproducible

Next, we evaluated the reproducibility of our library preparation and HTS methods by comparing read count data across replicates and between library formats. We compared replicate preparations of cDNA libraries from a single blood draw from each of two alpacas (named Marvel and Mikka) and found the replicates to be consistent (**fig. S14A**,**B**). As expected, the variability increased for less abundant nanobodies, but overall showed good reproducibility. Next, we pooled the cDNA samples from both alpacas, prepared a phage display library from the pool, and compared sequencing data from this library to the cDNA. The cDNA and phage libraries showed somewhat less concordance than the replicates, but overall indicated that most nanobodies were carried through the phage display library preparation process unaffected (**fig. S14C**). These results indicate that our library preparation and sequencing methods produce reliable results.

#### 2.2.3 Sampled nanobodies represent a large fraction of the total alpaca repertoire

Then, we evaluated the proportion of the alpaca immune repertoire that was covered by our cDNA sequencing experiments, using the same samples as **section 2.2.2**. We clustered the sequences using our recommended NanoMAP clustering method and analyzed the distribution of clonal family abundances with preseq [6]. Preseq uses these distributions to estimate the total size of the repertoire, including clonal families that were not observed. From these values, we calculated the fraction of the estimated total that went unobserved (**fig. S15A**,**B**) and the estimated total itself (**fig. S15C**,**D**). Interestingly, we found large variations between the two alpacas, with Marvel’s repertoire size and fraction unobserved much lower than Mikka’s. With our recommended clustering we estimate that our sequencing experiments captured 70-80% of the clonal families in Marvel’s repertoire, but only about 50-60% of Mikka’s. Although these results indicate that there are many more clonal families that we could collect data for, these families are likely to be low-abundance, as evidenced by the absence of high-abundance families among those that are observed only in one replicate (**fig. S15E**,**F**). The low-abundance of the clonal families observed in only one replicate suggests that they have not expanded in response to immunization, and therefore are unlikely to bind any of the targets, making them irrelevant to nanobody discovery. Thus, although we are not able to cover the complete repertoire, we were able to sample a significant fraction of it, which is likely to contain a substantial majority of the clonal families of interest.

### 2.3 NanoMAP clustering enables rapid selection of nanobody families with desired properties

After demonstrating that our clustering method was useful in generating accurate clonal families, we combined it with our phage display panning methods to determine how effectively and efficiently we could search the alpaca immune repertoire for nanobodies that had been elicited following immunization with known virulence factors and that showed promise as components of therapeutic agents. We selected virulence factor targets based on three major criteria: 1) prior conventional nanobody discovery work that we could compare with NanoMAP results, 2) the likely applicability of nanobodies in treating the associated disease, and 3) the global prevalence or importance of the disease.

#### 2.3.1 Characterization of the anti-schistosome virulence factor nanobody repertoire

The WHO estimates that about 600M people live in areas of the developing world endemic for schistosomes, parasitic worms that directly affect approximately 200M people worldwide [33]. We have reported on four surface proteins from Schistosoma mansoni (Sm) that are critical for the parasites’ survival in their mammalian hosts: 1) a tegumental acetylcholinesterase (SmTAChE) [29], 2) a phosphodiesterase/pyrophosphatase (SmNPP5) [3], 3) carbonic anhydrase (SmCA) [8], and 4) alkaline phosphatase (SmAP) [4]. We sought nanobodies binding these factors as research reagents and possible components of therapeutic or diagnostic agents. To efficiently identify nanobodies against all four targets, we immunized two alpacas with a pool of all four targets and produced a nanobody phage display library (**fig. 1A**). We then panned the library on each of the four targets individually, as well as on living larval Sm worms. After sequencing the library and the panned samples, we used NanoMAP to identify clonal families and calculate log_2_-fold-change (LFC) and GFold values for each family, yielding binding information for each family to each target. The binding phenotypes of the top-ranking clonal families to the four targets are shown in **fig. 4A**. These values allowed us to select the most enriched families for each target and filter out any families that showed nonspecific binding (e.g. binding to multiple targets; **section 4.7.1**), resulting in the identification of *>*30 families binding each target. Finally, the whole pathogen binding data allowed us to determine which nanobody families bind their targets at exposed epitopes on the parasite surface. Among the highest-ranked clonal families, most of those binding to SmNPP were also significantly enriched when panned on intact parasites. Thus, NanoMAP was able to simultaneously identify the most promising nanobody families targeting each virulence factor as well as those binding to epitopes on the living parasite surface.

**Figure 4:**
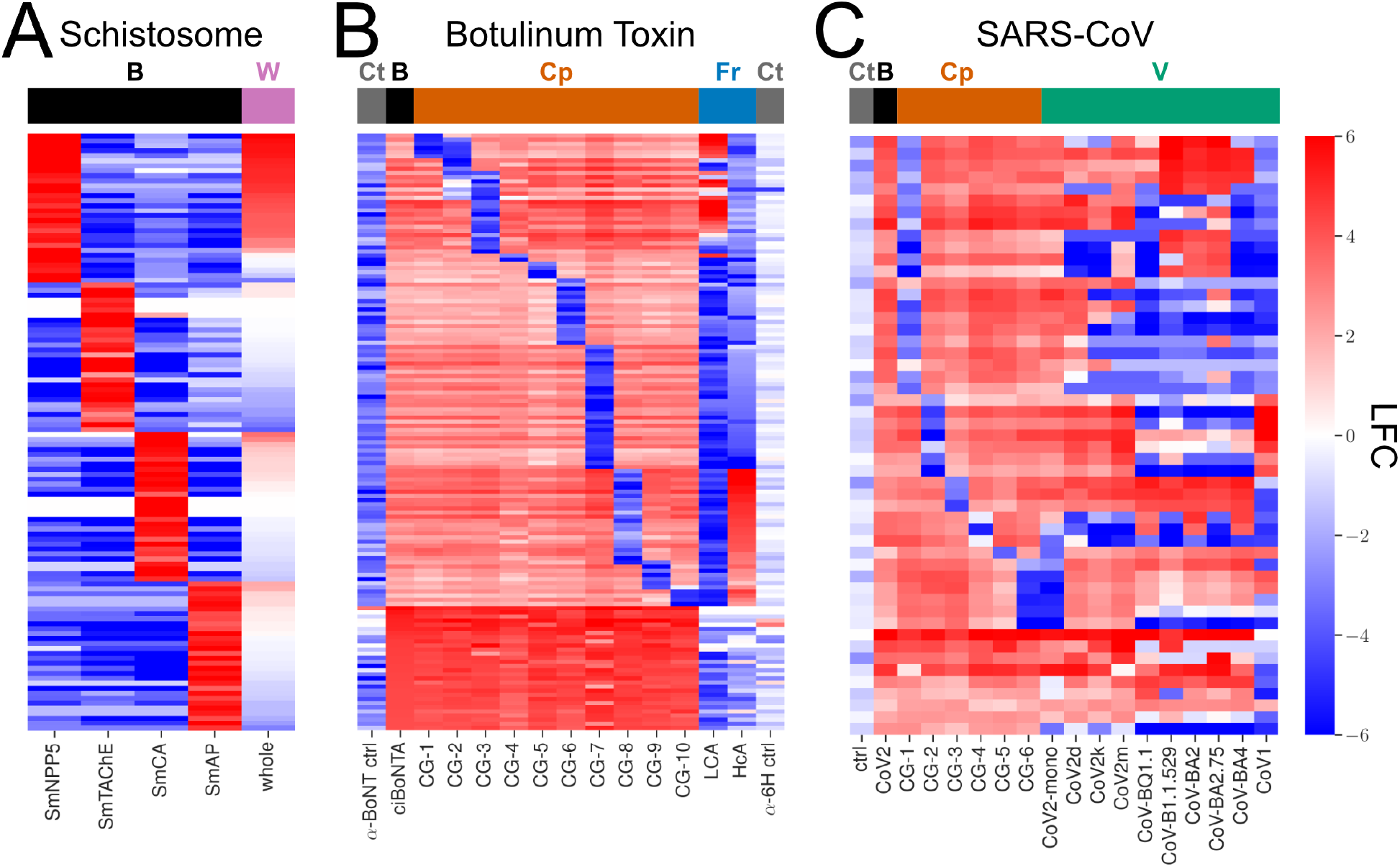
Panning experiments and meta-clustering facilitate identification of promising nanobody candidates. A-C) Heatmaps showing the LFC between the pre- and post-panning phage display library for selected clonal families, with high positive values denoting strong binding. A) LFC values for panning experiments on the four selected Schistosome surface enzymes: SmNPP5, SmTAChE, SmCA, and SmAP (black column label; basic panning). The final column indicates LFC values for panning experiments performed on whole schistosome parasites (pink column label; whole pathogen panning). B) LFC values for panning experiments on BoNT/A alone (black column label), in the presence of 10 different competitor nanobodies (orange column label; competitor panning), or on different domains of the toxin (blue column label; fragment panning). First and last columns show LFC values for a panning performed in the absence of BoNT/A (gray column label; control panning). C) LFC values for panning experiments with SARS-CoV2 spike protein alone (black column label), in the presence of 6 different competitors (orange column label), or using different variants of the spike protein (teal column label). CoV variants from throughout the COVID19 pandemic were selected in addition to a monomeric CoV2 spike, and an older variant (CoV1). The first column shows binding in the absence of any spike protein (gray column label).

#### 2.3.2 Characterization of the botulinum antitoxin nanobody repertoire

Botulinum neurotoxins (BoNTs) are CDC Category A biothreat agents and are responsible for a severe form of food poisoning. There are seven major BoNT serotypes, the most common and dangerous form is BoNT/A. We have previously characterized numerous anti-BoNT/A nanobodies recognizing a broad range of epitopes. These nanobodies can be divided into 10 “competition groups” (CGs) of nanobodies with overlapping epitopes identified through binding competition assays. Several of these CGs contain potent toxin neutralizing nanobodies, indicating that they bind near critical sites for BoNT function. The binding sites for many of these nanobodies have been characterized by X-ray crystallography studies [19, 18, 12, 37]. To broaden our options for nanobodies within the different CGs and identify nanobodies binding at new sites on the toxin, we panned our existing phage display library on a catalytically inactive BoNT/A holotoxin (ciBoNTA) in the absence or presence of competitor nanobodies representing each CG, and performed otherwise identical controls omitting ciBoNTA. We again used NanoMAP to define clonal families and calculate GFold and LFC values for each panning relative to the initial library. We then filtered for families that were highly enriched when panning on ciBoNTA alone, but were blocked from binding in the presence of each of the 10 CG representatives (**fig. 4B, section 4.7.2**). Using this approach, we readily “rediscovered” the nanobody families we had identified in prior studies [19, 18, 23] (see **fig. 5**), all of which were found in the CGs to which they had previously been experimentally assigned. We also identified many previously undiscovered families in each CG. Interestingly, some nanobody families were blocked by representatives from two different CGs, suggesting that the binding sites of the two CGs are adjacent, with the family’s epitope overlapping both. We also found many nanobody families that were not competed by any of our CG representatives (**fig. 4B**), providing an opportunity to identify nanobodies binding previously uncharacterized epitopes.

**Figure 5:**
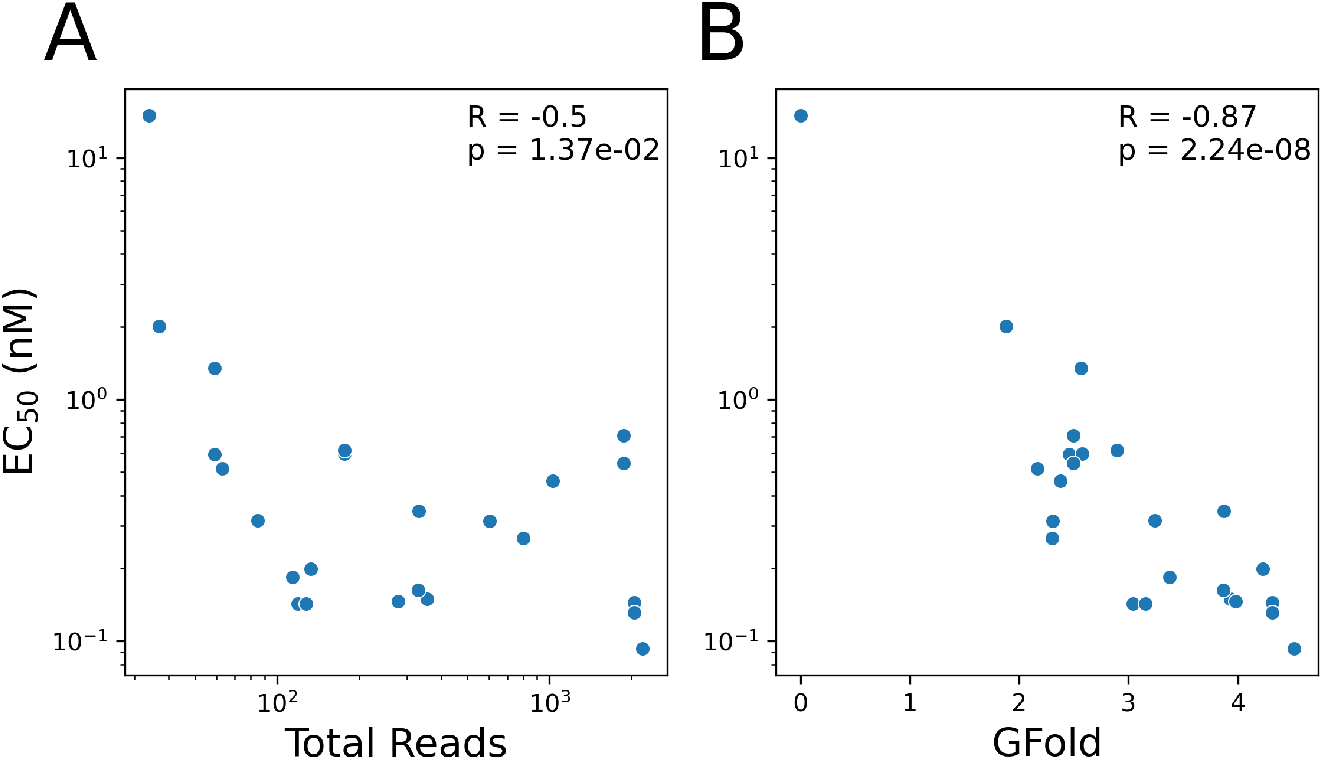
NanoMAP enrichment values correlate with binding strength. Scatter plots showing the correlation coefficient (R) and p-value (p) for the relationship between the EC_50_ for a each nanobody, measured by dilution ELISA, and values obtained from HTS experiments. A) Using the GFold value comparing the phage library abundance of the nanobody’s family to the abundance after basic ciBoNTA panning. B) Using the log_10_ of the total read counts for the family that the nanobody was part of in the ciBoNTA-panned sample.

In separate panning experiments performed in parallel with the competition pannings, we selected for nanobodies that bound to either of two BoNT/A domains, both of which are known to retain their functional conformations when expressed independently. Specifically, we panned on the BoNT/A heavy chain C-terminal domain (HcA; receptor binding domain) and the light chain (LCA; protease domain), and calculated GFold and LFC values for these experiments. As expected, there was a strong association between the competition and fragment binding data (**fig. 4B**). For example, families that were blocked by competitors representing CGs 8-10 clearly bound the receptor binding domain (HcA). All three representatives of these CGs have been shown to bind HcA, and two are known to block receptor binding, neutralizing toxicity [19]. Similarly, many members of CGs 1-4 bound the LCA domain. However, some did not, suggesting that the epitopes of these non-binding families overlap with other BoNT/A domains. Given that some families in CG2 are also in CG 1 or 3, it is likely that all three epitopes are very close to each other. Thus, the binding sites for those families that do not bind LCA alone must include regions of the LCA as well as nearby regions of other domains. The remaining competition groups must bind to epitopes that at least partially depend on the heavy chain N-terminal domain (HnA; transcytosis domain), located between LCA and HcA. The clear correlation between competition grouping and domain binding shows that both provide insight into the position of nanobody binding on the target. In total, these findings highlight the ability of NanoMAP to completely replicate our large body of preceding studies that used conventional methods to discover BoNT/A-binding nanobodies [23, 19, 18] and to simultaneously generate a more complete characterization of the anti-BoNT/A alpaca nanobody repertoire in a much shorter time frame.

#### 2.3.3 Discovery of SARS-CoV epitopes with broad recognition of natural variants

The spike protein of SARS-CoV viruses is the viral surface protein that interacts with the host ACE2 receptor during the infection process. Antibodies binding to some epitopes on the spike protein can prevent infection and this protein is routinely employed for CoV2 vaccination. As a result, we immunized alpacas with SARS-CoV2 spike (Wuhan) responsible for the COVID19 pandemic, prepared a nanobody phage display library and identified nanobodies that represented six different CGs. Similarly to the BoNT panning experiments, we used representatives of these six CGs to assign each member of our phage display library to a CG. In parallel, we panned our library on spike protein from several CoV2 variants that arose during the COVID19 pandemic, as well as to SARS-CoV1 (responsible for the 2002 SARS pandemic). Finally, we panned on monomeric CoV2 spike, to compare to the trimeric form used in the basic and competitor pannings. After processing the resulting data with NanoMAP and filtering for families with strong binding (**section 4.7.3**), we found at least one clonal family in each CG, as well as several families that were not part of any CG (**fig. 4C**). Notably, the different CGs display unique CoV1/2 variant specificities. Nanobodies in CG1 generally display highly specific binding to the original CoV2 spike, with a few exceptions. Nanobodies in CG2 bind to the early pandemic variants (*δ, κ*, and *μ*) and CoV1, but poorly bind the omicron variants. In contrast, nanobodies in CG3 generally bind well to all the CoV2 variants but not to CoV1 spike. Finally, CGs 5 and 6 appear to bind sites present only on trimeric CoV2 spike, but not on monomeric spike. These binding epitopes likely depend on a trimer-specific conformation or binding at an interface between subunits of the spike trimer. The CG5 and 6 epitopes appear to be much more conserved across CoV variants, allowing several clonal families identified in these CGs bind to all CoV2 variants tested, as well as CoV1. These results further demonstrate the ability of NanoMAP to generate rich binding information, in this case, allowing the selection of variant-agnostic clonal families whose binding is much less likely to be affected by viral evolution.

#### 2.3.4 NanoMAP results correlate with low-throughput nanobody characterization

Our results with schistosomes, BoNT, and CoV demonstrate NanoMAP’s ability to identify diverse binding phenotypes across a broad range of targets. Across the three datasets, NanoMAP is able to identify more highly enriched clonal families than other clustering methods without artificially breaking large families apart (**fig. S16**). To test whether these enrichment assessments (GFold) reflected individual nanobody binding affinity measurements, we evaluated whether the family-level GFold values derived from NanoMAP correlated with the measured binding strength of a member of the clonal family. We selected 24 nanobodies from our BoNT dataset, each from a distinct clonal family, spanning the range of non-negative GFold values observed in our basic ciBoNTA panning experiment. We measured the EC_50_ for each nanobody by performing dilution ELISAs using ciBoNTA capture conditions that closely matched those used in the panning study (**fig. S17A-C**). We then compared the EC_50_ values to the GFold and abundance values for each nanobody’s family (**fig. 5**). We found a strong (*R* = −0.87) and statistically significant (p = 2.2 · 10^−8^) correlation between EC_50_ and GFold, but a much weaker correlation with abundance in the ciBoNTA-panned library (*R* = −0.50). The large improvement in correlation when using GFold demonstrates the importance of fold-change based measures in the identification of high-affinity nanobody families; library screens that use abundance-based selection techniques are likely to miss out on strong but low-abundance candidates. These results also demonstrate that NanoMAP produces reliable binding values that are predictive of more direct experimental binding measurements.

## 3 Discussion

The results produced by NanoMAP demonstrate its ability to rapidly characterize alpaca immune repertoires and identify strong candidate nanobodies based on a variety of relevant properties including binding strength, variant specificity, and binding site. Our immunization and library preparation methods provide a reproducible and clean sample of a large fraction of each alpaca’s full immune repertoire. These comprehensive libraries can then be characterized through many parallel panning experiments which provide detailed data on the binding characteristics of each nanobody in the repertoire. Finally, NanoMAP’s clustering method groups the repertoire into clonal families with clear improvements in sequence and functional coherence over existing clustering methods. By aggregating data across related sequences, NanoMAP improves the reliability of the binding information provided for each nanobody family, resulting in enrichment values that are highly correlated with independent measurements of binding strength. Additionally, comparing NanoMAP data with results from previously published work employing conventional nanobody discovery methods [19, 18, 23] showed that NanoMAP predictions closely matched prior experimental data.

Critically, the success of our methods depends on accurate clustering; if a clonal family is “contaminated” with sequences from other clonal families having binding behaviors distinct from the correctly clustered family, family-level enrichment calculations would become less meaningful or even uninformative. Thus, by producing highly accurate clonal families, NanoMAP increases the signal-to-noise ratio for each enrichment value without compromising the quality of the results. To achieve the best results, we recommend the following: for datasets much larger than those presented here, SCOPer average-linkage clustering with a distance cutoff of 0.35 will likely produce good clusterings in a reasonable time frame and with reduced memory usage. However, for datasets of comparable or smaller size to ours, we recommend NanoMAP average-linkage individual clustering with a hierarchical distance cutoff of 0.467, and a merge distance cutoff of 0.25, as these settings are likely to produce a higher-quality clustering, at the cost of increased time and memory requirements. The quality of the clustering with these recommended parameters should be checked by computing silhouette scores, and if it is not satisfactory, the more computationally intensive NanoMAP meta-clustering method should be run.

When used together as described above, our immunization, panning, and clustering methods have the potential to greatly accelerate the rate of nanobody discovery by increasing the number of targets that can be used simultaneously, expanding the depth of characterization of immune repertoires, and providing more information about the phenotype of each clonal family. These rich characterizations are not limited to the types of panning assays performed here. Our methods could be readily adapted to identify nanobodies that block specific binding sites on the target by using mAbs with known epitopes, substrates, or binding partners of the target as competitors. These and other possible variations in panning conditions, such as pH or salinity changes, could be used to screen for a wide range of possible binding phenotypes tailored to fit the context of each target and the goals of the nanobody discovery campaign. Thus, our panning methods are readily adaptable to new targets and allow rapid identification of nanobodies with desirable binding properties against arbitrary targets.

In addition to these improvements to the nanobody discovery process, NanoMAP provides a starting point to more thoroughly characterize immune responses. NanoMAP could be paired with time series data of cDNA libraries collected over the course of sequential immunizations to understand how the dynamics of each clonal family relate to its affinity, improving our understanding of the affinity maturation process. It could also be paired with structure prediction methods to increase confidence when multiple members of a family or competition group are predicted to bind in the same location. Finally, NanoMAP could be used to augment training of deep learning models to predict nanobody binding either by providing family labels as additional input features, or by improving the quality of the binding data that the model is trained to predict. Overall, we expect the tools we have developed here to improve nanobody discovery directly and facilitate the development of new tools that further improve the discovery process.

## 4 Methods

### 4.1 Alpaca Immunization, Phage Display Library Preparation, and Panning

Alpaca immunization and nanobody-display phage library construction methods employed for these studies have been previously reported [16]. Schistosome proteins were all recombinant proteins, expressed in mammalian cells and shown to retain their enzymatic functions. The BoNT/A immunogen was the same as described in a previous publication [23]. The CoV immunogen was recombinant trimeric CoV2 spike protein (Wuhan; purchased from ACRO Biosystems). The three libraries employed for the schistosome, BoNT/A and CoV spike studies were produced from pools of 9.3 × 10^6^, 6.5 × 10^6^, and 1.5 × 10^7^ independent clones containing nanobody inserts respectively, and multiple aliquots were stored in 15% glycerol LBamp2%glucose at −80C. Nanobody-displaying phage preparation from thawed aliquots of the stored libraries and antigen panning were all generally performed as previously reported [16]. Specifically, 500 uL of 1 ug/mL of targets were coated to Nunc ImmunoTubes at 4°C overnight or captured to a tube previously coated with 500 uL of the target-capturing antibody and then washed and blocked overnight with 4%milk in PBS, 1% Tween20. For targets with hexahistidine tags (CoV, schistosome), the capture antibody was 2.5 ug/mL of an anti-His-tag mAb (Genscript A00186) incubated at 4°C overnight. For targets without hexahistidine (ciBoNTA), we employed 1.5 ug/ml of nanobody, JDA-D12 [17], which binds a unique BoNT/A site on the protease domain that does not compete with any known BoNT/A neutralizing nanobodies. After coating the capture antibody, we next incubated the tube in 500 ul containing 0.5 ug/ml of the target in 4%milk in PBS, 1% Tween20 for 1 hr at room temperature. After each step, tubes were washed three times with PBS, 0.1% Tween20 (PBS/Tween). To block phage binding to specific antigen binding sites, pannings were performed as above - but prior to phage addition, the antigens were pre-incubated for 30m with a high concentration of a previously characterized mAb or nanobody, typically at 50-100 ug/mL and this competitor solution remained throughout the subsequent target incubation. All pannings had a final volume 500 uL PBS/Tween with or without competitors. For live Sm larval worm panning, similar methods as above were used except that the live Sm (about 10,000 larvae) were in suspension, we omitted Tween in the buffers and employed mild centrifugation to wash the parasites (500 ×g for 2 min for each wash). To each tube, 100 uL of nanobody-displaying phage library containing 10^10^-10^11^ phage was added and incubated for 1h at room temperature (RT). Each tube was then washed 15 times with PBS/Tween at RT. Following these rapid washes, each tube was filled with PBS/Tween and incubated for one hour at 37°C on a rotator. This final wash was removed prior to phage elution. To elute the phage remaining in the washed tube, 500 uL of fresh ER2738 cells were added, incubated for 15 minutes at RT with rotation, and then removed to a second tube. Finally, 500 uL of 0.2M glycine pH 2.2 was added to each panning tube, incubated 10 minutes at RT with rotation, neutralized with 80 uL of 1M Tris pH 9 and combined with the ER2738 elutions as the final panning eluates for nanobody repertoire sequencing. An aliquot of the nanobody-display phage library prior to panning (termed “source”) was also saved for sequencing.

### 4.2 Phage Library Sequencing

In preparation for Illumina sequencing of the nanobody repertoires, the phage eluted in each panning eluate and the now infected ER2738 cells were added to 10 ml of SB with 100 ug/mL ampicillin, 10 ug/mL tetracycline and 2% glucose (SB/amp/tet) and incubated overnight at 250 RPM on a shaker at 37°C. Several 80 uL aliquots of the source libraries, equivalent to the amount used in each panning, were incubated in 1 mL of fresh overnight ER2738 cells, brought to 10 mL in SB/amp/tet and grown overnight as above. Phagemid preparations employing Promega miniprep kits were obtained from aliquots of each overnight culture. The nanobody repertoires of the source library and each panning eluate were separately amplified by PCR employing uniquely barcoded Illumina sequencing primer pair combinations. The sequences of the different barcoded primers employed are shown in **Table S1-2**. The 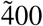 bp PCR products of each amplification were purified using the WizardR Plus SV minipreps DNA Purification System, quantified by Nanodrop, and equal amounts (by mass of DNA) of each product were pooled. The PCR pool was adjusted to 15 ug/mL and shipped to the Tufts University Core Facility (TUCF). Sequencing was performed on 100 uL of our PCR pool employing a MiSeqi100 with a 5M or 25M run type and PE300 read length (V3 600 cycles).

### 4.3 Sequence Assembly, Annotation, and Filtering

Sequences were processed using the pRESTO [35] and Change-O [13] packages from Immcantation to filter out low-quality reads and count duplicates. Then, NCBI IgBLAST [38] and ANARCI [9] were used to annotate the V/D/J and CRD regions of each sequence. Further details can be found in **section S1.1**.

### 4.4 Sequence (Meta-)clustering

Nanobody sequence diversity is generated in two stages, an initial V/D/J recombination event, and subsequent cycles of mutation and selection for improved antigen binding [24]. These steps naturally produce a clonal lineage where each surviving V/D/J recombination diversifies into a clonal family of related nanobody sequences that are highly likely to bind the same epitope of the same target. To infer clonal families from the HTS data, we clustered sequences into families that likely originated from the same V/D/J recombination event. We first grouped sequences with the same V and J segments and the same CDR lengths using Alakazam [13], split those groups into smaller sub-groups with similar sequences through hierarchical clustering using a distance metric similar to TCRdist [7]. Next, we took the most abundant sequence in each cluster as a representative and merged clusters based on an Leiden clustering of the representatives. Finally, to calculate the meta-clustering, we repeated the splitting and merging steps with different parameter settings, producing 150 distinct clusterings. Each of the clusterings was then scored, and the middle 60% of clusterings were used to calculate the final meta-clustering. Further details on the clustering methods and parameter choices are provided in the supplement (**section S1.2, table S3**).

### 4.5 Enrichment Calculations

Applying the methods described in **section 4.3** to the panning and sequencing experiments described in **section 4.2** results in a dataset that contains raw read counts of each observed DNA sequence in each of several experimental samples. These raw counts can be aggregated over amino acid sequences (grouping together different DNA sequences that encode the same amino acid sequence) or over clonal families determined by the clustering method described in **section 4.4**. We compared the read count values in samples generated from phage library panning experiments to those from the phage library prior to panning to calculate a quantitative measure of binding strength. We calculated two very similar values, LCF, and GFold, described in **sections S1.4.1 and S1.4.2**.

### 4.6 Evaluation of Clustering Methods

We evaluate NanoMAP’s clustering method against standard clustering methods in the field, namely SCOPer [25] and MMseqqs2 [30]. The first task is to identify optimal clustering parameters for each method. To identify these optimal clustering parameters, we selected three sequencing datasets, each targeting a separate pathogen - *Schistosoma mansoni* (schistosomes), *Clostridium botulinum* (Botulinum neurotoxin serotype A; BoNT/A), or SARS-CoV - and derived from immunization of a different pair of alpacas. The source nanobody display phage libraries from each alpaca pair had been previously subjected to conventional nanobody discovery from which numerous target binding nanobodies had been identified and characterized. The three selected datasets each contained sequences from the corresponding source library (**fig. 1A**; **section 4.1**) as well as enriched libraries created by panning the source library under various conditions (**fig. 1B**; **section 4.1**). The sequencing datasets contained roughly 4.0M, 14.1M, and 9.8M raw reads, resulting in a total of 195k, 582k, and 326k distinct amino acid sequences in the schistosome, BoNT/A, and SARS-CoV datasets, respectively, after quality control and filtering (**section 4.3**). All subsequent analysis is based on these three datasets.

#### 4.6.1 Clustering parameter sensitivity analysis

After filtering and annotating the sequences, we tested a broad range of parameter values for each clustering method. NanoMAP and SCOPer both use hierarchical clustering, and can therefor be run with single, average or complete linkage methods. MMseqs2 offers connected component and greedy set cover clustering options. We tested each of these linkage methods with a range of distance cutoffs, varying both the hierarchical and merging cutoffs for NanoMAP (**section S1.2.1**). Finally, we performed a hierarchical meta-clustering using the labels produced by the 150 individual NanoMAP clusterings, testing the same three linkage methods, and a range of distance cutoffs (**section S1.2.2**). We then evaluated the Silhouette index, phenotypic homogeneity, similarity to the “ground truth” clustering (ARI), and stability to data down-sampling of these clusterings as described below (**section 4.6**).

#### 4.6.1 Comparison to “ground truth”

To evaluate the quality of the clustering achieved by the methods described above, we established a “ground truth” clustering for a subset of sequences from each of three of our sequencing datasets. These sequences were selected to span a wide range of both CDR3 lengths and full-length sequence homologies but maintain a manageable size for manual annotation, resulting in 958, 1151, and 877 sequences selected from the Schistosome, BoNT/A, and SARS-CoV datasets, respectively.

To establish the “ground truth” clustering of these three sequence subsets, we produced a dendrogram based on a complete-linkage hierarchical clustering using the Levenshtein distance between full-length amino acid sequences. Using this dendrogram as a guide, we manually assigned sequences to clusters based on the similarity of their CDRs and whether they showed similar binding phenotypes. Sequences were split into a training set ( 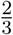 of the sequences) used for parameter optimization and a validation set (the remaining sequences) used for final performance evaluations such that each manually labeled cluster was fully contained in one of the two sets.

Each clustering was calculated using the full dataset from which the training and validation sequences were taken. Then, the “ground truth” clustering was used to evaluate computed clusterings by comparing the cluster labels for the subset of sequences in the training or validation set using the Adjusted Rand Index (ARI) [31, 15, 5]. ARI compares two clusterings based on how often they agree on whether a pair of sequences should be in the same or different clusters, and adjusts this value based on the expected value from a randomized clustering and the maximum possible value.

#### 4.6.3 Sequence silhouette index

To evaluate the overall homogeneity of our clusterings, we used an approximation of the Silhouette score [26]. The exact Silhouette score, *S*, is defined below for a clustering *C* with sequences 1 … *n*:

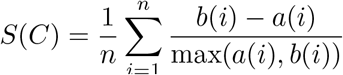

where *a*(*i*) is the average distance from sequence *i* to all other sequences in it’s cluster, and *b*(*i*) is the average distance from sequence *i* to all sequences in the nearest cluster that does not contain *i*. To avoid calculating all pairwise distances, we approximate *S* by assuming that all sequences in the same cluster have the same value of *a* and *b*. Then, for each cluster, *C*_1_ … *C*_*g*_ we approximate *a*(*C*_*g*_) by randomly sampling pairs within *C*_*g*_ and calculating the average of their CDR distance. For clusters where the number of possible pairs is small (*<* 1000), we calculate the average exactly. We approximate *b*(*C*_*g*_) by selecting the top 10 most abundant representatives from each cluster, and then using MMseqs2 to identify the top 10 closest matches (within the set of representatives) to each representative, excluding pairs within the same cluster. For each representative, the distance to each of the top 10 closest matches was calculated as in **section S1.2.1** and the lowest distance was recorded. The average of these distances across all representatives for each cluster is used to approximate *b*(*C*_*g*_). Finally, because our definition of CDR distance is constrained to the range [0, 1], we omit the normalization by max(*a*(*i*), *b*(*i*)) in order to create a global measure of cluster separation, rather than a local one.

#### 4.6.4 Phenotypic quality

To evaluate the ability of our clusterings to group sequences that share the same binding profile, we calculated the LFC values as described above (**section ??**) for each panning experiment in the dataset, and used these values to evaluate the phenotypic similarity within clusters. Specifically, for each sequence, we calculated the correlation distance (1 − *R*^2^) between its LFC values and the LFC values of each sequence in the dataset (note that for pairs of sequences with no variation in their LFC values, the correlation distance is undefined, and distances between such pairs are ignored in subsequent calculations). We then split these distances into two groups: distances to sequences in the same cluster as the sequence of interest (including the distance of zero to itself), and distances to all other sequences. We then used a standard one-sided MannWhitney U-test (assuming the within-cluster distances are smaller) to compare the two groups of distances, and collect the Z-statistic from this test for each sequence. For sequences in singleton clusters, there are not enough within-cluster distances to perform the test. In these cases we use a set of two zeros as the within-cluster distances. The score for the clustering as a whole is the average of the scores for each sequence in the dataset, divided by 50 to keep the scores in the range [0,1] for the clusterings we examine here.

#### 4.6.5 Down-sampling stability

To evaluate how clusterings are affected by changes in data sparsity, we clustered down-sampled versions of each dataset with varying degrees of down-sampling (see **section S1.6**) and evaluated the similarity of these clusterings to the clustering of the full dataset using the ARI. The stability of a clustering on a given dataset was defined as the lowest of these ARI values across 3-fold to 2187-fold down-sampling.

#### 4.6.6 Determination of optimal clustering parameters

To identify globally optimal clustering parameters, we first defined the composite score for a clustering of a dataset to be the product of the four scores outlined above for that clustering on that dataset. Then, for each dataset we applied a leave-one-out cross-validation approach, where optimal parameters for the dataset were selected to maximize the product of the composite scores of each of the other two datasets. The scores reported in **fig. 3** are the individual scores for the parameter settings identified by this method.

### 4.7 Selection of candidate nanobody clonal families

To select clonal families with the desired properties for each of the three groups of targets, we clustered the nanobody sequences separately for each target group using our optimal parameters (*d*_*a*_ = 0.467, *M* = average, and *d*_*merge*_ = 0.25). After aggregating read count data within each family, we calculated GFold and enrichment values comparing each panned sample to the phage library that was used as input to the panning experiments. We also calculated GFold values comparing each competitor panning to the relevant basic panning. GFold values were used to select clonal families with desirable properties as follows.

#### 4.7.1 Anti-schistosome virulence factor nanobodies

First, clonal families that showed non-specific binding were filtered out. Although this dataset did not contain a control panning, we were able to identify non-specific binding by finding clonal families that had GFold values above zero for more than one of the four schistosome surface enzyme targets. Since these enzymes are distinct in both sequence and structure, we expect that each nanobody would bind only one of the four. Thus, any clonal families with binding to more than one target were removed. Out of the remaining set of clonal families, we selected the families with the top 30 GFold values for each target, and removed any with Gfold values ≤ 0. The remaining set contains the strongest binding clonal families for each of the four targets.

#### 4.7.2 Anti-Botulinum neurotoxin serotype A nanobodies

First, clonal families displaying nonspecific binding were filtered out by removing any that had GFold *>* 0 for either of the controls. This set was further filtered to remove any clonal family with GFold ≤ 1 in the basic catalytically inactive BoNT/A (ciBoNTA) panning. For each competitor, we selected the 30 families with the lowest GFold scores in the presence of the competitor from the remaining set, removing any with competed GFold ≥ 0. Finally, we added the 30 families with the highest GFold in the basic panning out of the set that were not blocked by any competitor (and that met the initial filtering criteria). The resulting set was de-duplicated, resulting in the final set of families presented here, representing a range of different BoNT/A epitopes.

#### 4.7.3 Anti-SARS-CoV spike nanobodies

The CoV2 clonal families were selected in a similar manner to the BoNT/A families. Nonspecific binders were removed using the control panning by removing any families with Gfold *>* 0. From the resulting set we selected families with GFold *>* 1 in either the CoV2 or CoV1 panning, as long as they had GFold ≥ 0 for CoV2. As with the BoNT families, for each competitor, we selected the 30 families with the lowest GFold scores in the presence of the competitor from the remaining set, removing any with competed GFold ≥ 0. Finally, we added the set of families that were not blocked by any competitor (and that met the initial filtering criteria), and de-duplicated, resulting in a set of families recognizing diverse epitopes and variants of the CoV spike protein.

## Data and Code Availability

Raw sequencing data and processed data files including sequence, read count, LFC, GFold, and clustering information will be made publicly available upon request. Code for clustering and enrichment calculations, and to replicate the figures can be found at https://github.com/wlwhite-tufts/NanoMAP.

## Acknowledgements

We thank the Tufts BCB group and especially Faith Ocitti and Jocelyn Garcia for helpful discussions, and Lia Varghese for her illustrations. We thank Giselle Martinez, Marie Nguyen, and Field Willis for coming up with the name NanoMAP and designing the logo. We thank Lara Roche-Sudar for writing and figure design advice. We thank the Tufts University Core Facilities for the Illumina sequencing. This work was supported by NIH grants R01AI125704 (to C.B.S.) and 5K12GM133314-07 (supporting W.L.W.), as well as a seed grant from the Tufts Data Intensive Studies Center (Tufts DISC) (to L.J.C. and C.B.S.) as well as a Tufts Springboard grant (to C.B.S. and L.J.C.).

## References

[1] M. Arbabi-Ghahroudi. Camelid single-domain antibodies: promises and challenges as lifesaving treatments. International journal of molecular sciences, 23(9):5009, 2022.

[2] P. Bannas, J. Hambach, and F. Koch-Nolte. Nanobodies and nanobody-based human heavy chain antibodies as antitumor therapeutics. Frontiers in immunology, 8:1603, 2017.

[3] R. Bhardwaj, G. Krautz-Peterson, A. Da’dara, S. Tzipori, and P. J. Skelly. Tegumental phosphodiesterase SmNPP-5 is a virulence factor for schistosomes. Infection and Immunity, 79(10):4276–4284, 2011.

[4] R. Bhardwaj and P. J. Skelly. Characterization of schistosome tegumental alkaline phosphatase (SmAP). PLoS Neglected Tropical Diseases, 5(4):e1011, 2011.

[5] J. E. Chacón and A. I. Rastrojo. Minimum adjusted Rand index for two clusterings of a given size. Advances in Data Analysis and Classification, 17(1):125–133, 2023.

[6] T. Daley and A. D. Smith. Predicting the molecular complexity of sequencing libraries. Nature methods, 10(4):325–327, 2013.

[7] P. Dash, A. J. Fiore-Gartland, T. Hertz, G. C. Wang, S. Sharma, A. Souquette, J. C. Crawford, E. B. Clemens, T. H. Nguyen, K. Kedzierska, et al. Quantifiable predictive features define epitope-specific T cell receptor repertoires. Nature, 547(7661):89–93, 2017.

[8] A. A. Da’dara, A. Angeli, M. Ferraroni, C. T. Supuran, and P. J. Skelly. Crystal structure and chemical inhibition of essential schistosome host-interactive virulence factor carbonic anhydrase SmCA. Communications Biology, 2(1):333, 2019.

[9] J. Dunbar and C. M. Deane. ANARCI: antigen receptor numbering and receptor classification. Bioinformatics, 32(2):298–300, 2016.

[10] A. Evers, E. Guarnera, L. Pekar, and S. Zielonka. From discovery to the clinic: structural insights, engineering options, clinical, and ‘next wave’applications of camelid-derived single-domain antibodies. In mAbs, volume 17, page 2583210. Taylor & Francis, 2025.

[11] P. C. Fridy, M. P. Rout, and N. E. Ketaren. Nanobodies: from high-throughput identification to therapeutic development. Molecular & Cellular Proteomics, 23(12):100865, 2024.

[12] S. Gu, S. Rumpel, J. Zhou, J. Strotmeier, H. Bigalke, K. Perry, C. B. Shoemaker, A. Rummel, and R. Jin. Botulinum neurotoxin is shielded by NTNHA in an interlocked complex. Science, 335(6071):977–981, 2012.

[13] N. T. Gupta, J. A. Vander Heiden, M. Uduman, D. Gadala-Maria, G. Yaari, and S. H. Kleinstein. Change-O: a toolkit for analyzing large-scale B cell immunoglobulin repertoire sequencing data. Bioinformatics, 31(20):3356–3358, 2015.

[14] G. Houen. Therapeutic antibodies: an overview. Therapeutic Antibodies: Methods and Protocols, pages 1–25, 2021.

[15] L. Hubert and P. Arabie. Comparing partitions. Journal of classification, 2:193–218, 1985.

[16] J. J. Jaskiewicz, J. M. Tremblay, S. Tzipori, and C. B. Shoemaker. Identification and characterization of a new 34 kDa MORN motif-containing sporozoite surface-exposed protein, Cp-P34, unique to Cryptosporidium. International journal for parasitology, 51(9):761–775, 2021.

[17] C.-L. Kuo, G. A. Oyler, and C. B. Shoemaker. Accelerated neuronal cell recovery from Botulinum neurotoxin intoxication by targeted ubiquitination. PLoS One, 6(5):e20352, 2011.

[18] K.-h. Lam, J. M. Tremblay, K. Perry, K. Ichtchenko, C. B. Shoemaker, and R. Jin. Probing the structure and function of the protease domain of botulinum neurotoxins using single-domain antibodies. PLoS Pathogens, 18(1):e1010169, 2022.

[19] K.-h. Lam, J. M. Tremblay, E. Vazquez-Cintron, K. Perry, C. Ondeck, R. P. Webb, P. M. McNutt, C. B. Shoemaker, and R. Jin. Structural insights into rational design of single-domain antibody-based antitoxins against botulinum neurotoxins. Cell Reports, 30(8):2526–2539, 2020.

[20] R.-M. Lu, Y.-C. Hwang, I.-J. Liu, C.-C. Lee, H.-Z. Tsai, H.-J. Li, and H.-C. Wu. Development of therapeutic antibodies for the treatment of diseases. Journal of biomedical science, 27(1):1, 2020.

[21] F. D. Mast, P. C. Fridy, N. E. Ketaren, J. Wang, E. Y. Jacobs, J. P. Olivier, T. Sanyal, K. R. Molloy, F. Schmidt, M. Rutkowska, et al. Highly synergistic combinations of nanobodies that target SARS-CoV-2 and are resistant to escape. Elife, 10:e73027, 2021.

[22] J. Mukherjee, C. A. Ondeck, J. M. Tremblay, J. Archer, M. Debatis, A. Foss, J. Awata, J. H. Erasmus, P. M. McNutt, and C. B. Shoemaker. Intramuscular delivery of formulated RNA encoding six linked nanobodies is highly protective for exposures to three Botulinum neurotoxin serotypes. Scientific reports, 12(1):11664, 2022.

[23] J. Mukherjee, J. M. Tremblay, C. E. Leysath, K. Ofori, K. Baldwin, X. Feng, D. Bedenice, R. P. Webb, P. M. Wright, L. A. Smith, et al. A novel strategy for development of recombinant antitoxin therapeutics tested in a mouse botulism model. PloS one, 7(1):e29941, 2012.

[24] V. K. Nguyen, R. Hamers, L. Wyns, and S. Muyldermans. Camel heavy-chain antibodies: diverse germline VHH and specific mechanisms enlarge the antigen-binding repertoire. The EMBO journal, 2000.

[25] N. Nouri and S. H. Kleinstein. Somatic hypermutation analysis for improved identification of B cell clonal families from next-generation sequencing data. PLoS computational biology, 16(6):e1007977, 2020.

[26] P. J. Rousseeuw. Silhouettes: a graphical aid to the interpretation and validation of cluster analysis. Journal of computational and applied mathematics, 20:53–65, 1987.

[27] D. Saerens and S. Muyldermans. Single domain antibodies: methods and protocols, volume 911. Springer, 2012.

[28] S. Sharma, H. Byrne, and R. J. O’Kennedy. Antibodies and antibody-derived analytical biosensors. Essays in biochemistry, 60(1):9–18, 2016.

[29] P. Skelly and A. Da’dara. A novel, non-neuronal acetylcholinesterase of schistosome parasites is essential for definitive host infection. front immunol 14: 1056469. 10.3389/fimmu, 2023.

[30] M. Steinegger and J. Söding. Mmseqs2 enables sensitive protein sequence searching for the analysis of massive data sets. Nature biotechnology, 35(11):1026–1028, 2017.

[31] D. Steinley. Properties of the Hubert-Arable adjusted Rand index. Psychological methods, 9(3):386, 2004.

[32] A. Strehl and J. Ghosh. Cluster ensembles—a knowledge reuse framework for combining multiple partitions. Journal of machine learning research, 3(Dec):583–617, 2002.

[33] S. A.-L. Thétiot-Laurent, J. Boissier, A. Robert, and B. Meunier. Schistosomiasis chemotherapy. Angewandte Chemie International Edition, 52(31):7936–7956, 2013.

[34] H. Tsuruta, H. Yamazaki, R. Maeda, R. Tamura, J. Wei, Z. E. Mariet, P. Phloyphisut, H. Shimokawa, J. R. Ledsam, L. Colwell, et al. AVIDa-hIL6: a large-scale VHH dataset produced from an immunized alpaca for predicting antigen-antibody interactions. Advances in Neural Information Processing Systems, 36:42077–42096, 2023.

[35] J. Vander Heiden, G. Yaari, M. Uduman, J. Stern, K. O’Connor, D. Halfer, F. Vigneault, and S. Kleinstein. pRESTO: a toolkit for processing high-throughput sequencing raw reads of lymphocyte receptor repertoires (tech1p. 863). The Journal of Immunology, 192(Supplement 1):69–31, 2014.

[36] J. Xu, K. Xu, S. Jung, A. Conte, J. Lieberman, F. Muecksch, J. C. C. Lorenzi, S. Park, F. Schmidt, Z. Wang, et al. Nanobodies from camelid mice and llamas neutralize SARS-CoV-2 variants. Nature, 595(7866):278–282, 2021.

[37] G. Yao, K.-h. Lam, J. Weisemann, L. Peng, N. Krez, K. Perry, C. B. Shoemaker, M. Dong, A. Rummel, and R. Jin. A camelid single-domain antibody neutralizes botulinum neurotoxin A by blocking host receptor binding. Scientific Reports, 7(1):7438, 2017.

[38] J. Ye, N. Ma, T. L. Madden, and J. M. Ostell. IgBLAST: an immunoglobulin variable domain sequence analysis tool. Nucleic acids research, 41(W1):W34–W40, 2013.

[39] T. Yu, F. Zheng, W. He, S. Muyldermans, and Y. Wen. Single domain antibody: Development and application in biotechnology and biopharma. Immunological Reviews, 328(1):98–112, 2024.

