## Supplemental Information for "Comprehensive Mapping of Immune Nanobody Repertoires with NanoMAP"

#### S1 Supplemental methods

##### S1.1 Pre-processing details

To assemble, filter, and annotate the reads from our sequencing data, we used the procedure outlined below:

1. Paired end reads were assembled into full-length sequences (AssemblePairs; pRESTO).

2. Low-quality reads (phred quality, $Q < 20$) were removed (FilterSeq; pRESTO).

3. Reads with no identifiable forward or reverse primer sites were removed and primer sequences were trimmed off of the remaining sequences (MaskPrimers; pRESTO).

4. Reads were de-duplicated and counted (CollapseSeq; pRESTO).

5. V, D, and J segment assignments were made (igblastn; IgBLAST) for each sequence using a customized IgBLAST database (see **section S1.3**), and sequences without clear V or J assignments were removed (MakeDB; Change-O)

6. The start and end positions for translation were determined by sliding a "translation window" across the start or end of the DNA sequence, looking for the closest match to a predefined amino acid sequence. Sequences without a match were removed.

7. Translated sequences with an in-frame stop codon were removed.

8. Amino acid sequences observed only once in the dataset were removed.

9. The CDR amino acid sequences were extracted from each sequence using ANARCI.

The remaining amino acid sequences were used without further filtering in all subsequent steps.

##### S1.2 Meta-clustering details

###### S1.2.1 Initial clustering

Our initial clustering method proceeds through the three following steps, following a similar strategy to SCOPer [8] for the first two steps, and then performing a final merging step that allows groups with highly similar sequences, but different CDR lengths to be combined into more coherent groups.

**V/J/length grouping** First, we divided sequences into groups with matching V and J annotations and CDR lengths, using the groupGenes function in Alakazam [5]. Alakazam ignores D segment assignments because the high diversity of the surrounding region makes accurate D segment assignment difficult. Additionally, we defined a unique identifier for each combination of CDR1-3 lengths: $l = \sum_{i=1}^{3} 1000^{i-1} l_i$, where $l_i$ is the length of CDR$_i$, and used this value instead of CDR3 length as input to Alakazam to separate into groups with consistent length across all 3 CDRs.

**Hierarchical clustering** To produce high-resolution clusters within the V/J/length groups, we clustered each group above by hierarchical clustering using each linkage method, $M \in \{\text{single, average, complete}\}$, with a linkage-method-specific distance cutoff. We used 10 evenly-spaced values for each linkage method, with $d_s \in [0.05, 0.4]$, $d_a \in [0.05, 0.8]$, and $d_c \in [0.05, 0.95]$ for single, average, and complete linkages, respectively. Distances were defined using a custom CDR-centric distance function similar to TCRdist [2], defined below:

$$D_{CDR}(i,j) = \frac{\sum\limits_{k=1}^{3} D_H(s_i^k, s_j^k) w_k}{\sum\limits_{k=1}^{3} l_k w_k}$$

where $D_H$ is the non-normalized Hamming distance, $s_i^k$ is the sequence of the $k^{th}$ CDR of sequence $i$, and $l_k$ is the length of the the $k^{th}$ CDR. Note that due to the CDR length grouping, $l_k$ is the same for each $(i, j)$ pair.

**Cluster Merging** In order to allow for some flexibility in CDR lengths, clusters produced by the hierarchical clustering step above were merged together based on the similarity of a representative selected from each cluster. Specifically, the most abundant member of each cluster was selected as its representative, and used to construct a graph with edges connecting any representatives with CDR distance $< d_{merge} \in \{0.1, 0.15, 0.2, 0.25\}$. We then used Leiden community detection [11] to define communities in this graph, and merged any clusters from the hierarchical clustering step whose representatives were in the same community. In order to accommodate possible differences in CDR lengths, we adjusted out CDR distance definition as follows:

$$D_{CDR}(i,j) = \begin{cases} \frac{\sum_{k=1}^{3} L(s_i^k, s_j^k) w_k}{\sum_{k=1}^{3} \max(l_i^k, l_j^k) w_k} & \text{if } \max\limits_k |l_i^k - l_j^k| \leq \Delta l \\ 1 & \text{otherwise} \end{cases}$$

and $s_i^k$ and $l_i^k$ are the sequence and length of the $k^{th}$ CDR of sequence $i$, respectively, $L$ is the Levenshtein distance, $w_k > 0$ is the per-amino-acid weight of the $k^{th}$ CDR (we used 1, 1, and 3 for CDR1, CDR2, and CDR3 respectively; **table S3**), and $\Delta l$ is the maximum allowed difference in CDR length (we used 3).

##### S1.2.2 Sequence Meta-Clustering

To achieve a robust clustering across varied dataset compositions and sizes, we calculated many initial clusterings, and used the cluster labels from each to calculate a final meta-clustering. To generate diverse initial clusterings, we varied $M$, $d_s$, $d_a$, $d_c$, and $d_{merge}$, with all other parameters held constant. Parameter values and variations are noted above, and complied in **table S3**.

**Scoring** The resulting initial clusterings were scored using a metric inspired by the Silhouette Index [9], but much faster to calculate for large datasets. We defined the Fast Silhouette Index, $S_{fast}$, for clustering $C$ and sequences 1 ... $n$:

$$S_{fast}(C) = b_{avg}(C) - a_{avg}(C)$$

where $b_{avg}$ is the average between-cluster distance:

$$\frac{\sum_{i=1}^{n} \sum_{j \notin C(i)} d_{CDR}(i,j)}{\sum_{i=1}^{n} \sum_{j \notin C(i)} 1}$$

and $a_{avg}$ is the average within-cluster distance:

$$\frac{\sum_{i=1}^{n} \sum_{j \in C(i), i \neq j} d_{CDR}(i,j)}{\sum_{i=1}^{n} \sum_{j \in C(i), i \neq j} 1}$$

and $C(i)$ is the set of sequences in the same cluster as $i$.
To make the calculation of $S_{fast}$ actually fast, we approximated it by randomly sampling 2 sets of 10,000 pairs of sequences, one where both sequences in each pair are in the same cluster, and the other where they are in different clusters. We then calculate the average distance across pairs in each set and subtract them as described above to approximate $S_{fast}$.

**Meta-clustering** After scoring each individual clustering (including the hierarchical clusterigns, and all merged variants thereof), we retained clusterings with score quantiles $q_{low} < q < q_{high}$. We selected $q_{low} = 0.2$ and $q_{high} = 0.8$ because we found that the lowest scoring clusterings often contained clusters that were too broad while the highest scoring clusterings contained clusters that were too narrow. After individual clusterings were selected, meta-clustering was performed by hierarchical clustering with a distance cutoff of $d_{meta} \in (0, 1)$ and a linkage method of $M_{meta} \in \{\text{single, average, complete}\}$, using the Hamming distance between vectors of cluster labels (the fraction of non-matching labels) as the distance function. We tested many combinations of meta-clustering parameters to determine the parameters that perform the best across datasets and are the most stable across dataset sizes (**fig. 3, fig. S1,S5-7**).

#### S1.3 Addition of new V and J segments to IgBLAST database

To prevent exclusion of certain group of nanobody sequences from our analysis, we added several V and J segments matching the 5' and 3' ends of those sequences to the IgBLAST database. We first constructed a phylogenetic tree of all examples of this group that we had observed in prior work using the EMBL Clustal$\Omega$ web server [7], and selected several representatives from distinct regions of the tree. To obtain the appropriate ranges of nucleotide positions to include in the V an J segment, we aligned the full DNA sequences of several selected representatives from the excluded group to the existing V or J segments in the database and selected the range of positions that was covered in the alignment. We then translated the segments and used the IMGT alignment web app [3] to insert gaps into the newly identified V and J segments. Finally, we reverse-translated the gapped sequences, preserving the gaps and original codon usage and used the gapped DNA sequences to identify the boundaries of the CDRs and FWRs for entry into the alpaca.ndm.imgt (for V segments) and alpaca_gl.aux (for J segments) files. Finally, we added our gapped DNA sequences to the fasta files containing the V and J segment sequences and re-created the IgBLAST database using the modified files. New V segments were named IGHV5S1*01 ... IGHV5S1*07 and new J segments were named IGHJ8*01 ... IGHJ8*04 so that they could be easily distinguished from existing V and J segments whose numbering stops at IGHV4 and IGHJ7, respectively. The updated database was used for annotation and filtering of all sequences presented here.

#### S1.4 Enrichment calculation details

##### S1.4.1 log-Fold-Change (LFC)

For a set of $n$ amino acid sequences (or clonal families) with read count data for each of two experimental samples (usually before and after a panning experiment), $k_1^b$ ... $k_n^b$ and $k_1^a$ ... $k_n^a$:

$$\mathrm{LFC}_i = \log_2 \frac{\frac{k_i^a}{K^a} + \epsilon}{\frac{k_i^b}{K^b} + \epsilon}$$

where

$$K^x = \sum_{i=1}^{n} k_i^x \qquad \text{and} \qquad \epsilon = \frac{1}{\max(K^a, K^b)}$$

The constant $\epsilon$ is added to avoid division by zero or evaluating $\log_2(0)$.

##### S1.4.2 GFold

Although the LFC provides a direct assessment of enrichment of sequences (or clonal families), it is subject to significant experimental noise, particularly for sequences with low read counts, because of the quantized nature of read count data. To account for this uncertainty, we used GFold [4] to adjust LFC values based on estimations of the variance of the measurement. This results in the most confident and strong binding sequences (or clonal families) being ranked highest, and assigns a score of 0 to sequences with very low signal-to-noise ratios, allowing low-quality LFC values to be filtered out easily. We used $\alpha = 0.01$ in our GFold calculations, meaning that the true LFC value is expected to be closer to zero than the GFold score only $1\%$ of the time.

#### S1.5 Comparator clustering methods

**MMseqs** As a baseline clustering comparison, we clustered the same sequences with MMseqs2 [10] using a range of distance cutoffs, and two different clustering methods. MMseqs2 allows users to choose between two relevant clustering modes: greedy set cover (clusters groups of sequences that are all within the specified distance of a central sequence) and connected component (clusters groups of sequences where each sequence is within the specified distance of at least one other member of the group). We applied each of these methods to cluster all sequences in the dataset using only their CDR3 sequence.

**SCOPer** As an immune-receptor-specific clustering comparison, we clustered the same sequences with the hierarchicalClones function in SCOPer [8], using a range of distance cutoffs, in combination with each of the three linkage method options (single, average, and complete). We used ”nt” mode, meaning that sequences were clustered based on CDR3 nucleotide sequences.

#### S1.6 Dataset down-sampling

To produce smaller datasets to use in assessing the stability of different clustering methods to variations in dataset size (**fig. S2-12**), we used a semi-randomized approach intended to mimic an experiment with a smaller sequencing depth. Starting with a complete dataset (with sequences $1...n$ observed $c_1...c_n$ times) and a down-sampling factor, $\phi \in (0, 1]$, we generated an expected number of read counts for each sequence, $e_i = \phi \times c_i$. For each sequence, we then drew a down-sampled number of observations from a Poisson distribution and removed the sequence from the down-sampled dataset if the sampled value was zero. Finally, any sequences present in the "ground truth" clustering for the dataset that were removed by this process were added back in in order to allow a fair comparison across all values of $\phi$. We used $\phi = 3^{-k}$ with $k \in \{1, 2, 3, 4, 5, 6, 7\}$ as well as the full dataset (rather than sampling with $\phi = 1$).

#### S1.7 Error rate estimation

To estimate the error rate in our sequencing experiments, we performed PCR on a single plasmid containing a nanobody coding DNA using our standard PCR conditions. For each unique DNA sequence observed in the resulting dataset, we counted the number of times it was observed in the dataset, and aligned it to the expected sequence and tallied the number of mutations, insertions, and deletions and their positions relative to the expected sequence (with insertions and deletions marked as occurring at the position immediately preceding their location. Our overall per-read error rate was calculated as:

$$\varepsilon = \frac{\sum_{i=1}^{n} m_i c_i}{\sum_{i=1}^{n} c_i}$$

where $m_i$ is the total mutations, insertions, and deletions in sequence $i$, and $c_i$ is the number of times sequence $i$ was observed in the dataset.

#### S1.8 cDNA replicates

To assess the reproducibility of our sequencing results, we prepared cDNA from two different PBL pools ($2 \times 10^7$ cells each) that had been purified from the same 100 mL blood samples obtained from each of two alpacas (Marvel and Mikka) and used them to generate a nanobody-display phage library. We then performed both long hinge and short hinge PCRs on each cDNA sample with uniquely barcoded primers, purified the nanobody coding DNAs and pooled equal amounts of each for Illumina sequencing. Sequencing data from this pool were used in **figures S14-15**.

#### S1.9 Repertoire size estimation

To estimate the size of the alpaca nanobody repertoire, we used NanoMap to identify clonal families in the samples described in **section S1.8**, with a modified filtering step (**section S1.1**, step 8) that retains all amino acid sequences, not just those that are observed more than once. Retaining the singleton sequences is critical for accurate estimations of repertoire size. For each replicate sample, as well as pooled samples within alpacas, we used preseq [1] to estimate the true size of the repertoire based on the distribution of family (or individual nanobody) abundances. We repeated this process using individual nanobody counts or clonal family counts (clustering using our optimal NanoMAP parameters).

#### S1.10 Dilution ELISAs

Standard target capture dilution ELISAs were performed as previously described [6] employing ELISA plates pre-coated with 1.5 mg/ml JDA-D12 nanobody followed by the capture of 0.5 mg/ml of the ciBoNTA target. These ELISA conditions were chosen to mimic the conditions used in the panning experiment that generated the corresponding ciBoNTA enrichment data.

#### S1.11 Candidate family grouping analysis

Related families are identified by finding the connected components of a graph with families as nodes, and edges connecting similar families. Families are considered similar if they meet both of the following criteria:
1. The CDR distance between the representatives (most abundant sequence) of the families is $< 0.3$
2. The correlation coefficient calculated by comparing the enrichment values for the families across all panning experiments is $> 0.75$

#### S2 Supplemental figures and tables

**Supplementary Table 1: Barcoding primers for cDNA.** This table shows the sequences of the barcoded primers that were used to prepare the cDNA libraries for sequencing.

| Name | barcode | index | sequence |
|---|---|---|---|
| VHH-L1 | CTCAGTTC | i5 | AATGATACGGCGACCACCGAGATCTACACCGCTCAGTTCTCGTCGGCAGC GTCAGATGTGTATAAGAGACAGGGTGGTCCTGGCTGC |
| VHH-L2 | TAGATCGC | i5 | AATGATACGGCGACCACCGAGATCTACACTAGATCGCTCGTCGGCAGCGT CAGATGTGTATAAGAGACAGGGTGGTCCTGGCTGC |
| VHH-L3 | CTCTCTAT | i5 | AATGATACGGCGACCACCGAGATCTACACCTCTCTATTCGTCGGCAGCGT CAGATGTGTATAAGAGACAGGGTGGTCCTGGCTGC |
| VHH-L4 | TATCCTCT | i5 | AATGATACGGCGACCACCGAGATCTACACTATCCTCTTCGTCGGCAGCGT CAGATGTGTATAAGAGACAGGGTGGTCCTGGCTGC |
| VHH-L5 | AAGGAGTA | i5 | AATGATACGGCGACCACCGAGATCTACACAAGGAGTATCGTCGGCAGCGT CAGATGTGTATAAGAGACAGGGTGGTCCTGGCTGC |
| VHH-L6 | CTAAGCCT | i5 | AATGATACGGCGACCACCGAGATCTACACCTAAGCCTTCGTCGGCAGCGT CAGATGTGTATAAGAGACAGGGTGGTCCTGGCTGC |
| VHH-L7 | CGTCTAAT | i5 | AATGATACGGCGACCACCGAGATCTACACCGTCTAATTCGTCGGCAGCGT CAGATGTGTATAAGAGACAGGGTGGTCCTGGCTGC |
| VHH-L8 | GCGTAAGA | i5 | AATGATACGGCGACCACCGAGATCTACACGCGTAAGATCGTCGGCAGCGT CAGATGTGTATAAGAGACAGGGTGGTCCTGGCTGC |
| VHH-sh2 | TAAGGCGA | i7 | CAAGCAGAAGACGGCATACGAGATGTTCGCCTTAGTGACTGGAGTTCAGA CGTGTGCTCTTCCGATCTCTTCGCTGTGGTGCGC |
| VHH-sh3 | CGTACTAG | i7 | CAAGCAGAAGACGGCATACGAGATGTCTAGTACGGTGACTGGAGTTCAGA CGTGTGCTCTTCCGATCTCTTCGCTGTGGTGCGC |
| VHH-sh4 | AGGCAGAA | i7 | CAAGCAGAAGACGGCATACGAGATGTTTCTGCCTGTGACTGGAGTTCAGA CGTGTGCTCTTCCGATCTCTTCGCTGTGGTGCGC |
| VHH-sh5 | CGAGGCTG | i7 | CAAGCAGAAGACGGCATACGAGATGTCAGCCTCGGTGACTGGAGTTCAGA CGTGTGCTCTTCCGATCTCTTCGCTGTGGTGCGC |
| VHH-sh6 | AAGAGGCA | i7 | CAAGCAGAAGACGGCATACGAGATGTTGCCTCTTGTGACTGGAGTTCAGA CGTGTGCTCTTCCGATCTCTTCGCTGTGGTGCGC |
| VHH-sh7 | GTAGAGGA | i7 | CAAGCAGAAGACGGCATACGAGATGTTCCTCTACGTGACTGGAGTTCAGA CGTGTGCTCTTCCGATCTCTTCGCTGTGGTGCGC |
| VHH-lh2 | TCCTGAGC | i7 | CAAGCAGAAGACGGCATACGAGATGTGCTCAGGAGTGACTGGAGTTCAGA CGTGTGCTCTTCCGATCTGGTTTTGGTGTCTTGGG |
| VHH-lh3 | GGACTCCT | i7 | CAAGCAGAAGACGGCATACGAGATGTAGGAGTCCGTGACTGGAGTTCAGA CGTGTGCTCTTCCGATCTGGTTTTGGTGTCTTGGG |
| VHH-lh4 | TAGGCATG | i7 | CAAGCAGAAGACGGCATACGAGATGTCATGCCTAGTGACTGGAGTTCAGA CGTGTGCTCTTCCGATCTGGTTTTGGTGTCTTGGG |
| VHH-lh5 | GCTCATGA | i7 | CAAGCAGAAGACGGCATACGAGATGTTCATGAGCGTGACTGGAGTTCAGA CGTGTGCTCTTCCGATCTGGTTTTGGTGTCTTGGG |
| VHH-lh6 | ATCTCAGG | i7 | CAAGCAGAAGACGGCATACGAGATGTCCTGAGATGTGACTGGAGTTCAGA CGTGTGCTCTTCCGATCTGGTTTTGGTGTCTTGGG |
| VHH-lh7 | ACTCGCTA | i7 | CAAGCAGAAGACGGCATACGAGATGTTAGCGAGTGTGACTGGAGTTCAGA CGTGTGCTCTTCCGATCTGGTTTTGGTGTCTTGGG |

**Supplementary Table 2: Barcoding primers for phage plasmids.** This table shows the sequences of the barcoded primers that were used to prepare the phage display and panned libraries for sequencing.

| Name | barcode | index | sequence |
|---|---|---|---|
| JSC-F1 | ATGAGACG | i5 | AATGATACGGCGACCACCGAGATCTACACATATGAGACGTCGTCGGCAGC GTCAGATGTGTATAAGAGACAGCCCAACCGGCCATGGC |
| JSC-F2 | AGAGTAGA | i5 | AATGATACGGCGACCACCGAGATCTACACAGAGTAGATCGTCGGCAGCGT CAGATGTGTATAAGAGACAGCCCAACCGGCCATGGC |
| JSC-F3 | GTAAGGAG | i5 | AATGATACGGCGACCACCGAGATCTACACGTAAGGAGTCGTCGGCAGCGT CAGATGTGTATAAGAGACAGCCCAACCGGCCATGGC |
| JSC-F4 | ACTGCATA | i5 | AATGATACGGCGACCACCGAGATCTACACACTGCATATCGTCGGCAGCGT CAGATGTGTATAAGAGACAGCCCAACCGGCCATGGC |
| JSC-F5 | TCTCTCCG | i5 | AATGATACGGCGACCACCGAGATCTACACTCTCTCCGTCGTCGGCAGCGT CAGATGTGTATAAGAGACAGCCCAACCGGCCATGGC |
| JSC-F6 | TCGACTAG | i5 | AATGATACGGCGACCACCGAGATCTACACTCGACTAGTCGTCGGCAGCGT CAGATGTGTATAAGAGACAGCCCAACCGGCCATGGC |
| JSC-F7 | TTCTAGCT | i5 | AATGATACGGCGACCACCGAGATCTACACTTCTAGCTTCGTCGGCAGCGT CAGATGTGTATAAGAGACAGCCCAACCGGCCATGGC |
| JSC-F8 | CCTAGAGT | i5 | AATGATACGGCGACCACCGAGATCTACACCCTAGAGTTCGTCGGCAGCGT CAGATGTGTATAAGAGACAGCCCAACCGGCCATGGC |
| JSC-F9 | CTATTAAG | i5 | AATGATACGGCGACCACCGAGATCTACACCTATTAAGTCGTCGGCAGCGT CAGATGTGTATAAGAGACAGCCCAACCGGCCATGGC |
| JSC-sh2 | CTCTCTAC | i7 | CAAGCAGAAGACGGCATACGAGATGTGTAGAGAGGTGACTGGAGTTCAGA CGTGTGCTCTTCCGATCTCTTCGCTGTGGTGCGC |
| JSC-sh3 | CAGAGAGG | i7 | CAAGCAGAAGACGGCATACGAGATGTCCTCTCTGGTGACTGGAGTTCAGA CGTGTGCTCTTCCGATCTCTTCGCTGTGGTGCGC |
| JSC-sh4 | GCTACGCT | i7 | CAAGCAGAAGACGGCATACGAGATGTAGCGTAGCGTGACTGGAGTTCAGA CGTGTGCTCTTCCGATCTCTTCGCTGTGGTGCGC |
| JSC-sh5 | GGAGCTAC | i7 | CAAGCAGAAGACGGCATACGAGATGTGTAGCTCCGTGACTGGAGTTCAGA CGTGTGCTCTTCCGATCTCTTCGCTGTGGTGCGC |
| JSC-sh6 | GCGTAGTA | i7 | CAAGCAGAAGACGGCATACGAGATGTTACTACGCGTGACTGGAGTTCAGA CGTGTGCTCTTCCGATCTCTTCGCTGTGGTGCGC |
| JSC-sh7 | CGGAGCCT | i7 | CAAGCAGAAGACGGCATACGAGATGTAGGCTCCGGTGACTGGAGTTCAGA CGTGTGCTCTTCCGATCTCTTCGCTGTGGTGCGC |
| JSC-sh8 | TACGCTGC | i7 | CAAGCAGAAGACGGCATACGAGATGTGCAGCGTAGTGACTGGAGTTCAGA CGTGTGCTCTTCCGATCTCTTCGCTGTGGTGCGC |
| JSC-sh9 | ATGCGCAG | i7 | CAAGCAGAAGACGGCATACGAGATGTCTGCGCATGTGACTGGAGTTCAGA CGTGTGCTCTTCCGATCTCTTCGCTGTGGTGCGC |
| JSC-lh2 | TAGCGCTC | i7 | CAAGCAGAAGACGGCATACGAGATGTGAGCGCTAGTGACTGGAGTTCAGA CGTGTGCTCTTCCGATCTGGTTTTGGTGTCTTGGG |
| JSC-lh3 | ACTGAGCG | i7 | CAAGCAGAAGACGGCATACGAGATGTCGCTCAGTGTGACTGGAGTTCAGA CGTGTGCTCTTCCGATCTGGTTTTGGTGTCTTGGG |
| JSC-lh4 | CCTAAGAC | i7 | CAAGCAGAAGACGGCATACGAGATGTGTCTTAGGGTGACTGGAGTTCAGA CGTGTGCTCTTCCGATCTGGTTTTGGTGTCTTGGG |
| JSC-lh5 | CGATCAGT | i7 | CAAGCAGAAGACGGCATACGAGATGTACTGATCGGTGACTGGAGTTCAGA CGTGTGCTCTTCCGATCTGGTTTTGGTGTCTTGGG |
| JSC-lh6 | TGCAGCTA | i7 | CAAGCAGAAGACGGCATACGAGATGTTAGCTGCAGTGACTGGAGTTCAGA CGTGTGCTCTTCCGATCTGGTTTTGGTGTCTTGGG |
| JSC-lh7 | TCGACGTC | i7 | CAAGCAGAAGACGGCATACGAGATGTGACGTCGAGTGACTGGAGTTCAGA CGTGTGCTCTTCCGATCTGGTTTTGGTGTCTTGGG |
| JSC-lh8 | AATTCTGC | i7 | CAAGCAGAAGACGGCATACGAGATGTGCAGAATTGTGACTGGAGTTCAGA CGTGTGCTCTTCCGATCTGGTTTTGGTGTCTTGGG |
| JSC-lh9 | GGCCTCAT | i7 | CAAGCAGAAGACGGCATACGAGATGTATGAGGCCGTGACTGGAGTTCAGA CGTGTGCTCTTCCGATCTGGTTTTGGTGTCTTGGG |

**Supplementary Table 3: Parameter settings for initial clustering.** This table details the parameter settings or ranges of values that were used to produce the initial clusterings used in meta-clustering. The names of the steps match the headings in **figure 2**, and the meaning of the parameters is described in **section S1.2.1**. Parameters that were varied are listed with values $> 1$ in the 'number of values' column, with values sampled evenly across the corresponding range.

| Step Name | Parameter | Range | Number of Values | Description |
|---|---|---|---|---|
| hierarchical | $M$ | single<br>average<br>complete | 3 | Linkage method for hierarchical clustering. |
| hierarchical | $d_s$ | 0.05 - 0.4 | 10 | Distance cutoff for single-linkage hierarchical clustering. |
| hierarchical | $d_a$ | 0.05 - 0.8 | 10 | Distance cutoff for average-linkage hierarchical clustering. |
| hierarchical | $d_c$ | 0.05 - 0.95 | 10 | Distance cutoff for complete-linkage hierarchical clustering. |
| hierarchical & merge | $w_1, w_2$ | 1 | 1 | Per-position weight for CDR1,CDR2. |
| hierarchical & merge | $w_3$ | 3 | 1 | Per-position weight for CDR3. |
| merge | $d_{merge}$ | 0.1 - 0.25 | 4 | Distance cutoff for edges in graph of cluster representatives. |
| merge | $\Delta l$ | 3 | 1 | Maximum allowed CDR length difference. |

**Supplementary Table 4: Dataset size and clustering runtimes.** Table showing the dataset and cumpute requirement information. Dataset size is measured by the number of reads collected across all samples (Reads), the number of distinct nanobody amino acid sequences identified after QC and filtering (Nanobodies), and the number of clonal families (Families) identified by each method using optimal parameters. Each clustering method is evaluated based on the wall-clock runtime (Runtime), maximum memory usage (Memory), and number of CPU cores used (CPUs) when clustering each of the three datasets.

| Dataset | Method | Reads | Nanobodies | Families | Runtime | Memory | CPUs |
|---|---|---|---|---|---|---|---|
| Schisto | NanoMAP (ind) | 4,009,051 | 194,868 | 24,023 | 00:26:37 | 8.05 GB | 10 |
| Schisto | NanoMAP (meta) | 4,009,051 | 194,868 | 21,963 | 01:12:04 | 21.20 GB | 10 |
| Schisto | SCOPer | 4,009,051 | 194,868 | 18,958 | 00:15:08 | 3.33 GB | 1 |
| Schisto | MMseqs2 | 4,009,051 | 194,868 | 21,030 | 00:00:32 | 74.82 MB | 1 |
| BoNT/A | NanoMAP (ind) | 14,078,864 | 582,166 | 34,533 | 01:14:32 | 24.76 GB | 10 |
| BoNT/A | NanoMAP (meta) | 14,078,864 | 582,166 | 32,471 | 02:48:24 | 49.52 GB | 10 |
| BoNT/A | SCOPer | 14,078,864 | 582,166 | 27,666 | 01:10:27 | 12.22 GB | 1 |
| BoNT/A | MMseqs2 | 14,078,864 | 582,166 | 54,867 | 00:00:36 | 114.37 MB | 1 |
| CoV | NanoMAP (ind) | 9,817,998 | 325,591 | 20,457 | 00:43:34 | 23.46 GB | 10 |
| CoV | NanoMAP (meta) | 9,817,998 | 325,591 | 19,229 | 01:34:26 | 40.13 GB | 10 |
| CoV | SCOPer | 9,817,998 | 325,591 | 17,670 | 00:24:12 | 9.22 GB | 1 |
| CoV | MMseqs2 | 9,817,998 | 325,591 | 58,862 | 00:00:32 | 94.07 MB | 1 |

**Supplementary Table 5: Nanobody sequences used in BoNT/A ELISAs** This table contains the sequences and $EC_{50}$ values of the 24 nanobodies used in the dilution ELISAs measuring $EC_{50}$ for binding to ciBoNTA (**fig. 5, S17**). $EC_{50}$ values are reported in nM.

| Name | $EC_{50}$ | Sequence |
|---|---|---|
| JDQ-A5 | 14.97 | LVQPGGSLRLSCAASAGNLDYYAIGWFRQAPGKEREGVSCISSSDGSTVYTDSVKGRFTISRDNTKNTVD<br>LQMDNLKPEDTAVYYCATVVNYYCTAGGSIHASPYEIWGQGTQVTVSS |
| JDQ-B5 | 0.59 | LVQDGGSLRLSCTTSGSIDNFNAIEWYRQAPGKQRELVASISSDGRRTNYADSVKGRFTISGDNAKNTVY<br>LQMNSLKPEDTAVYYCHRPFTHYWGEGTQVTISS |
| BXAJ-1 | 0.15 | LVQPGGSLTLSCAASGSIFEIYAMGWFRQAPGKQRELVATITSDGHTNTADSVLGRFTISRDSAKNTVFL<br>RMNSLKPEDTAVYFCNAQGRITVATMRPQYEYWGQGTQVTVSS |
| BXAJ-2 | 0.60 | SVQAGGSLRLTCVHSGTMFMLKAMGWARQAPGKQRESVATITTDGHIDYADSVKGRFTISRDNAKNTVYL<br>QMNSLNLEDTAVYYCNADYGVPHYWGQGIQVTVSS |
| BXAJ-3 | 0.32 | LVQPGGSLRLSCAASGLTMDYYEIGWFRQAPGKEREGVSSIRSHDSFTYYADSVKGRFTVSRDNAKNTVF<br>LQMNSLKPEDTAIYYCAVDVTPYYAGSYLDAADYDYWGQGTQVTVSS |
| BXAJ-4 | 0.55 | LVQPGGSLRLSCAASGPTYDIGWFRQAPGKERETVSCMTSSDGSTYYADSAKGRFTISRDIAKNTVYLEM<br>NSLKPEDTAVYYCAADLTGTGSDYDDACGFDAWGQGTQVTVSS |
| BXAJ-5 | 0.62 | LVQTGGSLRLSCTSSESIFSRTTMEWYRQPPGKQREWVASMPSDGRTWYSHSVEGRFTFSRDDAQDTLYL<br>QMNSLKPEDTAVYYCHSDLGDLWGQGTQVTVSA |
| BXAJ-6 | 2.00 | LVQAGGSLRLSCAASGSFSSINTMGWYRQTPGVQLDLVASITSSGTTDTAGSVKGRFIISRDNAKNTVYL<br>QMNSLKPEDTDVYYCYARVSPPLDGDYLLDEYEYYGQGTQVTVSS |
| BXAJ-7 | 1.35 | LVQAGDSLRLSCAASSTLFDSDTIGWYRQPPGKQRQLIAVLRSGNTGDYNDSIKDRFTISRDNAKNTVYL<br>QMNDLKPEDTAVYTCALENRYSRQRNSWGQGTQVTVSS |
| BXAJ-8 | 0.14 | LVQAGGSLRLSCAASGDTICISTMGWYRQAPGKERELVATVTGGGDIFYADSVKGRFTVSRDNAKNSVYL<br>QMDSLRPEDTAVYYCYADSNWCDSFGTYDFWGQGTQVTVSS |
| BXAJ-9 | 0.35 | LVQTGGSLRLSCAASGTMNDLNPMAWYRQGPGKQRESVAFISSTGITKYGDSVKGRFTISRDGAKNTVYL<br>QMNSLKPEDTGVYYCNIVDRSKDYWGQGTQVTVSS |
| JDQ-F12 | 0.27 | LVQPGGSLRLSCAASGFAVGSRYMSWVRQAPGKGLEWVSSIEPDPGPTWDADSVKGRFTISRDDAKNTLY<br>LQMSNLQPEDTGVYYCATGYRTNTRLPGGSWGQGTQVTVSS |
| JDQ-G5 | 0.14 | LVQPGGSLRLSCTASGITNGRFVMAWYRQTPGNEREFVASISSSGGTTSYAPAVKGRFTISRDNNAENAI<br>ILQMNSLKPEDTAMYYCRTPNYWGQGTQVTVSS |
| BXAJ-10 | 0.16 | WVQPGGSLKLSCAASGSISSINGMGWYRQAPGKQRELVATITRAGFTNYLDSVNRRFTISRDNAKNTLYL<br>QMNSLKPEDTAVYYCNAQMGDYYSGPIVDYWGQGTQVTVSS |
| BXAJ-11 | 0.52 | LVQPGGSLRLSCAASGFAFSSSAMTWVRQAPGKGLEWVSTIYSGGHTRTYADSVNGRFTISRDDAKNTLY<br>LQMNNLQPEDSAVYYCANKGGQYPDYAPLSGQGTRVTVSS |
| BXAJ-12 | 0.14 | FVQAGGSLRLSCAASGNIDSIKGMGWYRQAPGKQRELVATITQRDFTNYADSVMGRFTISRENAEKVFLQ<br>MNSLKPEDTAVYYCHAWSGTPFETAYREYWGQGTQLTVSS |
| BXAJ-13 | 0.20 | LVQPGGSLRLSCAVSGTISRDDILGWYRQAPGRQRELVADIRSPGVETYASSLQGRFTISRDKDENTVYL<br>QMSDLKPEDTAVYYCNSRSIFDRSAGYWGQGTQVTVSS |
| BXAJ-14 | 0.13 | LVQPGGSLRLSCTASGNINNINGMGWYRQAPGKQRELVATITDRGFTNTADSVVGRFTISREDTTKVFLQ<br>MTSLKPEDTAVYFCNAWSGRPFAPDWREYWGQGTQLTVSS |
| BXAJ-15 | 0.15 | FVQAGGSLRLSCVASGSSWTFDVMGWYRQAPGKQRELVATITRDKFTNYESSVKDRFTISGDTAKNTVYL<br>QMNALKPEDTAVYYCHGQNRPFFGQPEYWGQGTQVTVSS |
| JDQ-H7 | 0.09 | LVQVGGSLRLSCVVSGSDISGIAMGWYRQAPGKRREMVADIFSGGSTDYAGSVKGRFTISRDNAKKTSYL<br>QMNNVKPEDTGVYYCRLYGSGDYWGQGTQVTVSS |
| XCL-A3 | 0.46 | LVQPGGSLRLSCAANGDTLEHDTLGWFRQAPGKEREAVSCISSADDGTYYADSVKGRFTISRDNAKGTVD<br>LRMNNLKPEDTAVYYCATAGVPSSGSGCYYLRYDLWGQGTQVTVSS |
| XCL-C5 | 0.71 | LVQPGGSLRLSCTVTEFTRDYYDIGWFRQAPGKGRETVSCMTISDYSTYYADSVKGRFTISRDNAKNTVY<br>LQMNSLKPEDTSTYYCAVDLTGGRGSDYEDACGFDYWGQGTQVTVSS |
| XCL-E11 | 0.31 | LVEAGGSLRLSCTASGRFFRANTMGWYRQAAGKERELVAVMINAGATDGVINYADSVKGRFTISRDNAKS<br>MVYLQMNSLKPEDTAVYYCYARTYVAGWGQGTQVTVAS |
| XCL-F10 | 0.18 | LVQAGGSLRLSCEASGTISPRNAMGWYRQAPGKQRELVATETNAGFKNYADSVKGRFTVSRDNAKNAVNL<br>QMNNLRPDDTAVYYCNIDFPPWGREIWGQGTQVTVSS |

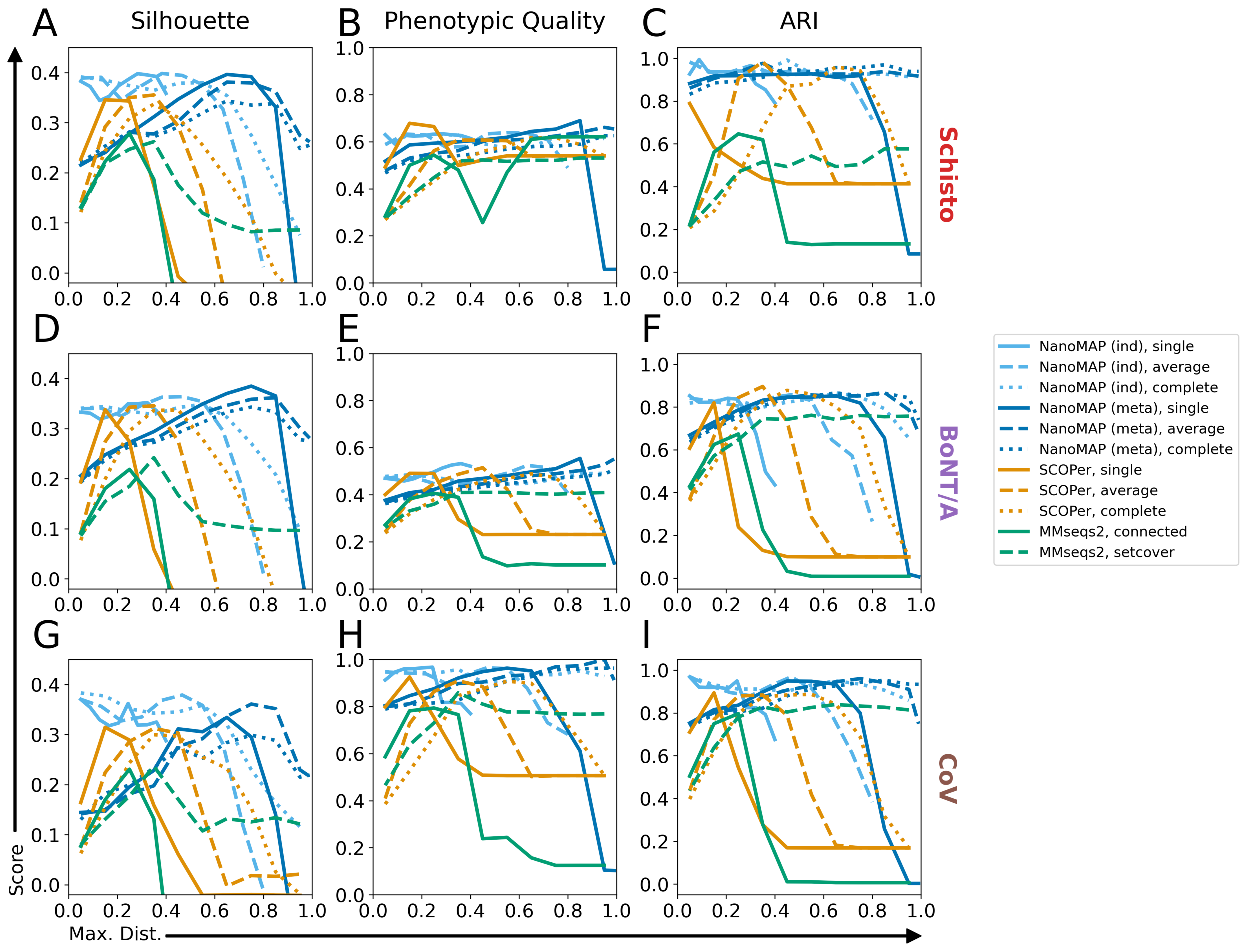


**Supplementary Figure 1: Clustering quality across methods and distance cutoffs.** Plots showing how variations in clustering distance cutoff impact the Silhouette index (A,D,G), phenotypic quality (B,E,H), and ARI (C,F,I) in each of our three datasets, when using NanoMAP individual (light blue), ManoMAP meta- (dark blue), SCOPer (orange), or MMseqs2 (green) clustering. For NanoMAP and SCOPer clusterings, solid, dashed and dotted lines indicate single, average, and compete linkage methods, respectively. For individual clusterings, $d_s$, $d_a$, or $d_c$ was varied with $d_{merge}$ held constant at 0.25.

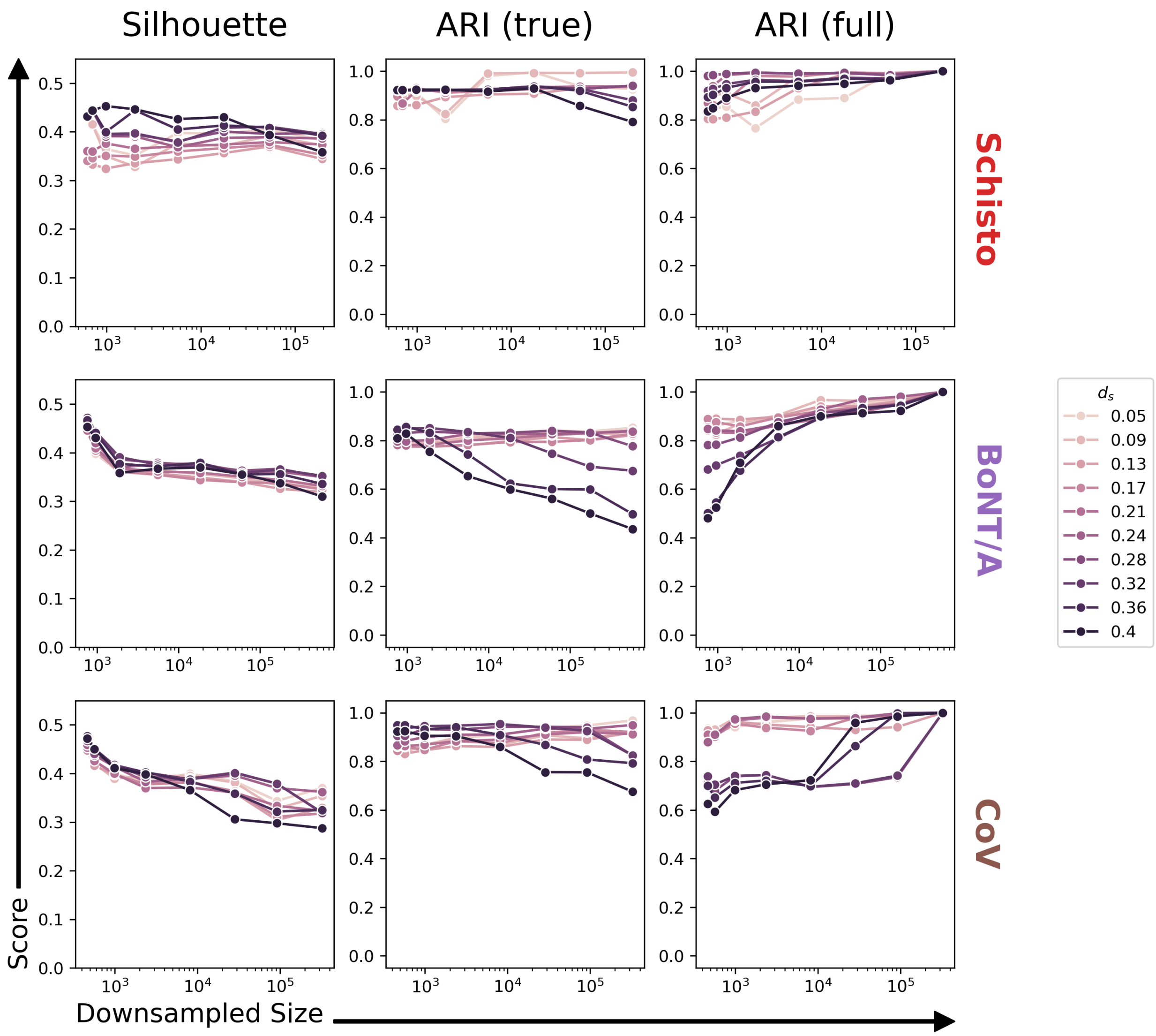


**Supplementary Figure 2: Sensitivity of single-linkage individual clustering to dataset size.** Plots showing the relationship between dataset size and the cluster quality metrics described in **section 4.6** for individual with $M$ = single, and $d_{merge} = 0.25$, with varying values of $d_s$. Smaller datasets were obtained by randomly removing individual reads from the full-size dataset. Down-sampled size indicates the number of distinct amino acid sequences present in the down-sampled dataset after filtering out sequences with only one read. ARI (true) indicates the ARI resulting from the comparison to the "ground truth" clusters as in **figure 3**, while ARI (full) indicates comparison to the clustering of the full-sized dataset using the same parameters.

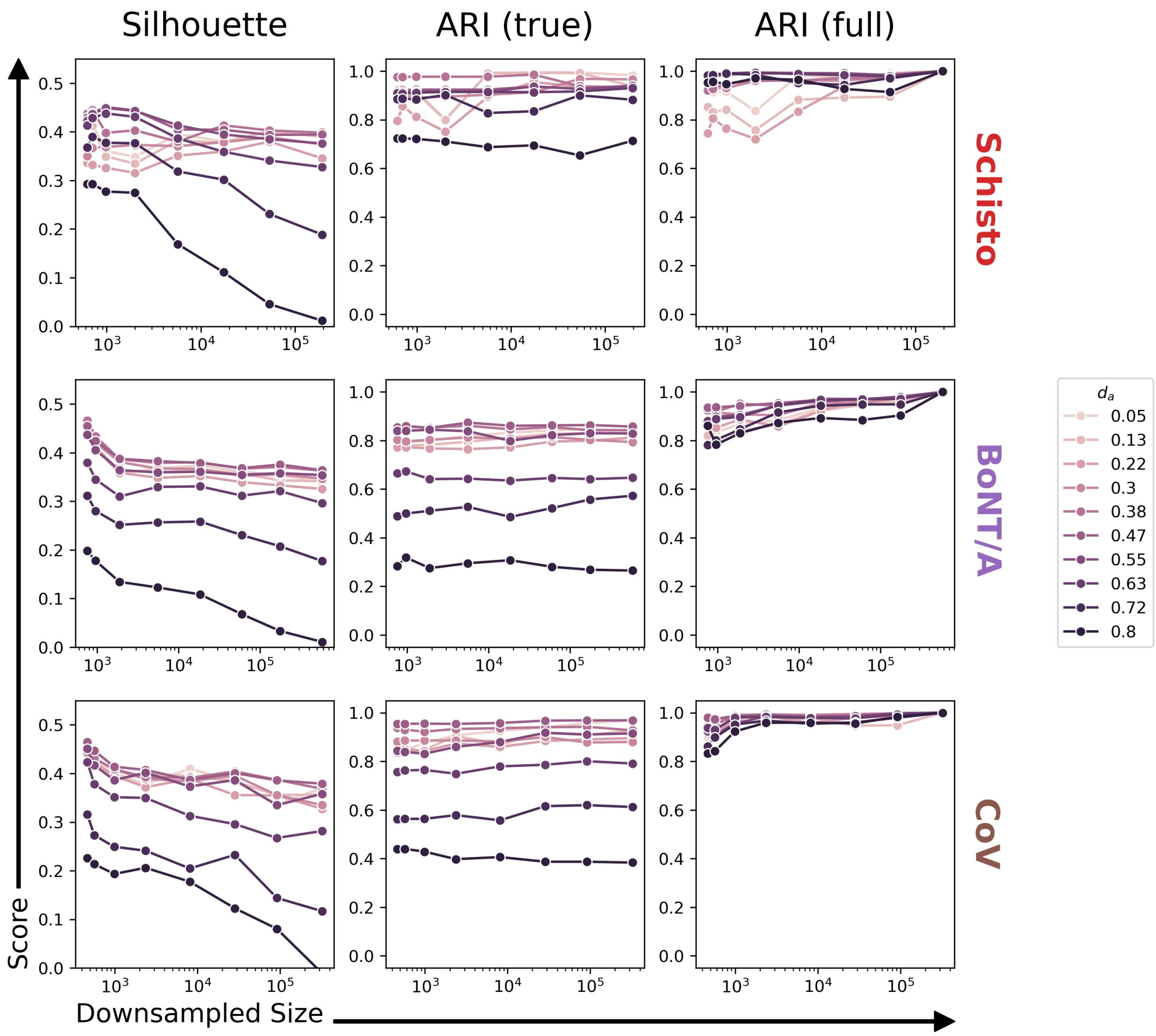


**Supplementary Figure 3: Sensitivity of average-linkage individual clustering to dataset size.** Plots as in **figure S2** showing results for $M =$ average, and $d_{merge} = 0.25$, with varying values of $d_a$.

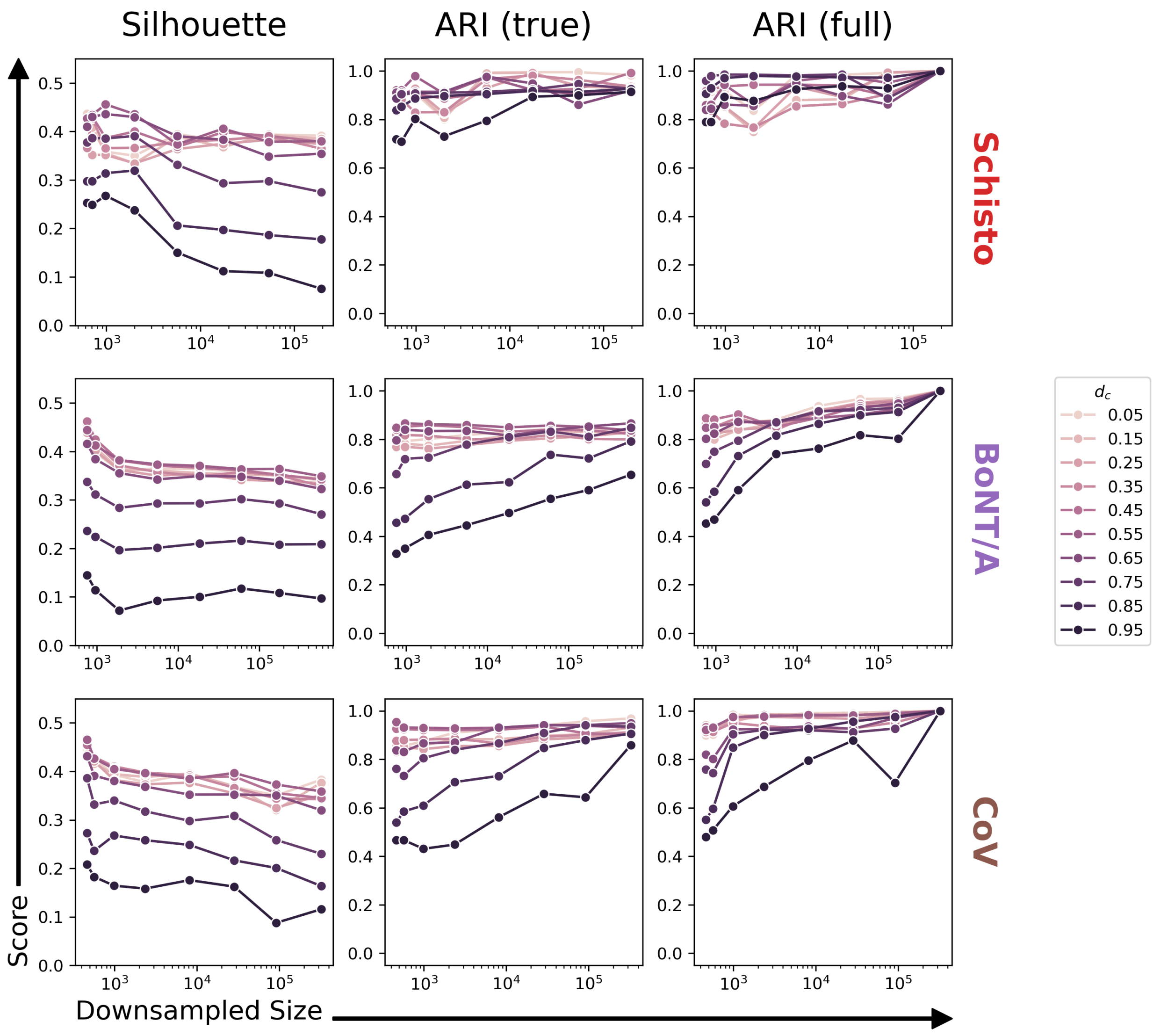


**Supplementary Figure 4: Sensitivity of complete-linkage individual clustering to dataset size.** Plots as in **figure S2** showing results for $M =$ complete, and $d_{merge} = 0.25$, with varying values of $d_c$.

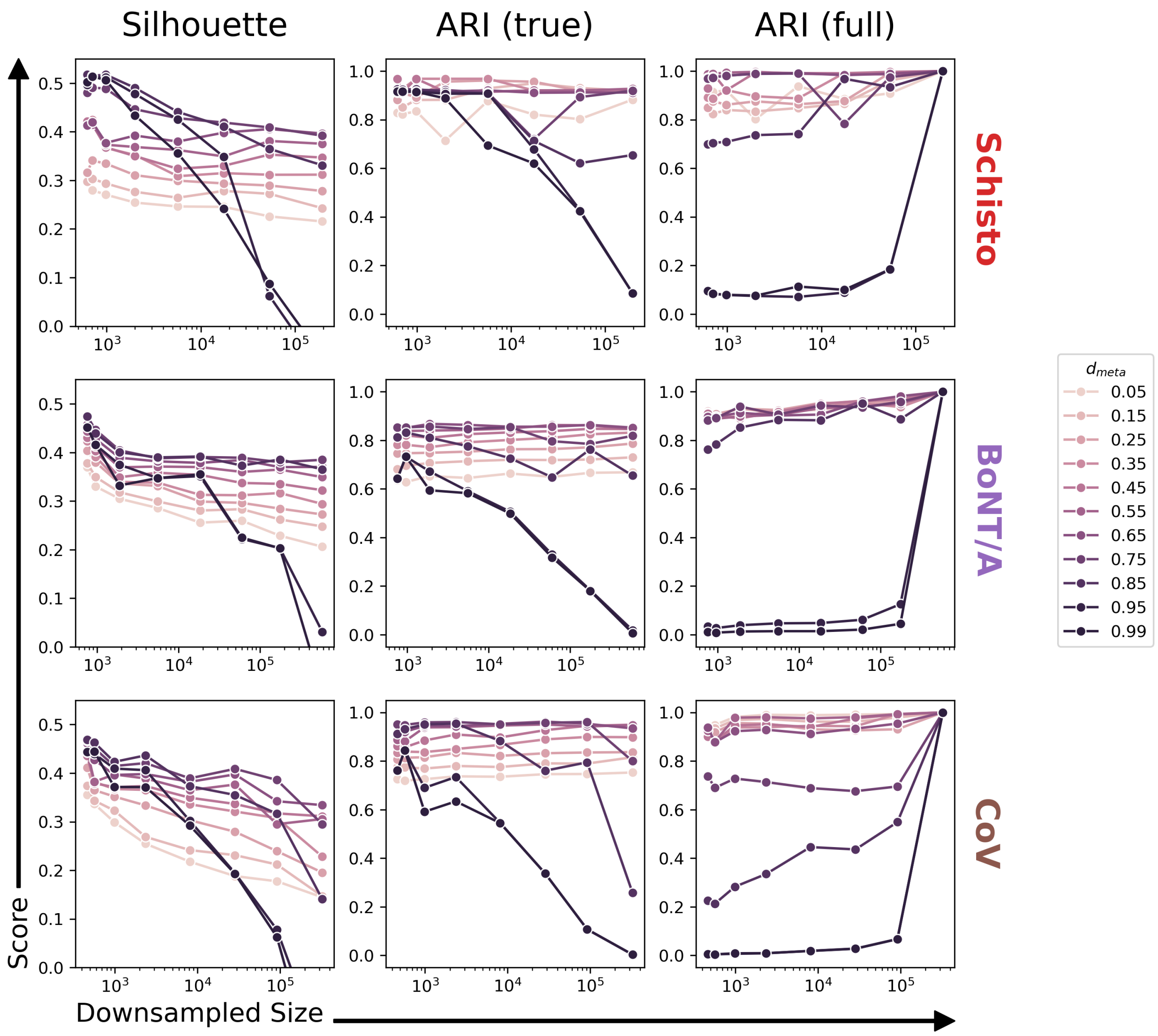


**Supplementary Figure 5: Sensitivity of single-linkage meta-clustering to dataset size.** Plots as in **figure S2** showing results for meta-clustering with $M_{meta}$ = single, and varying values of $d_{meta}$.

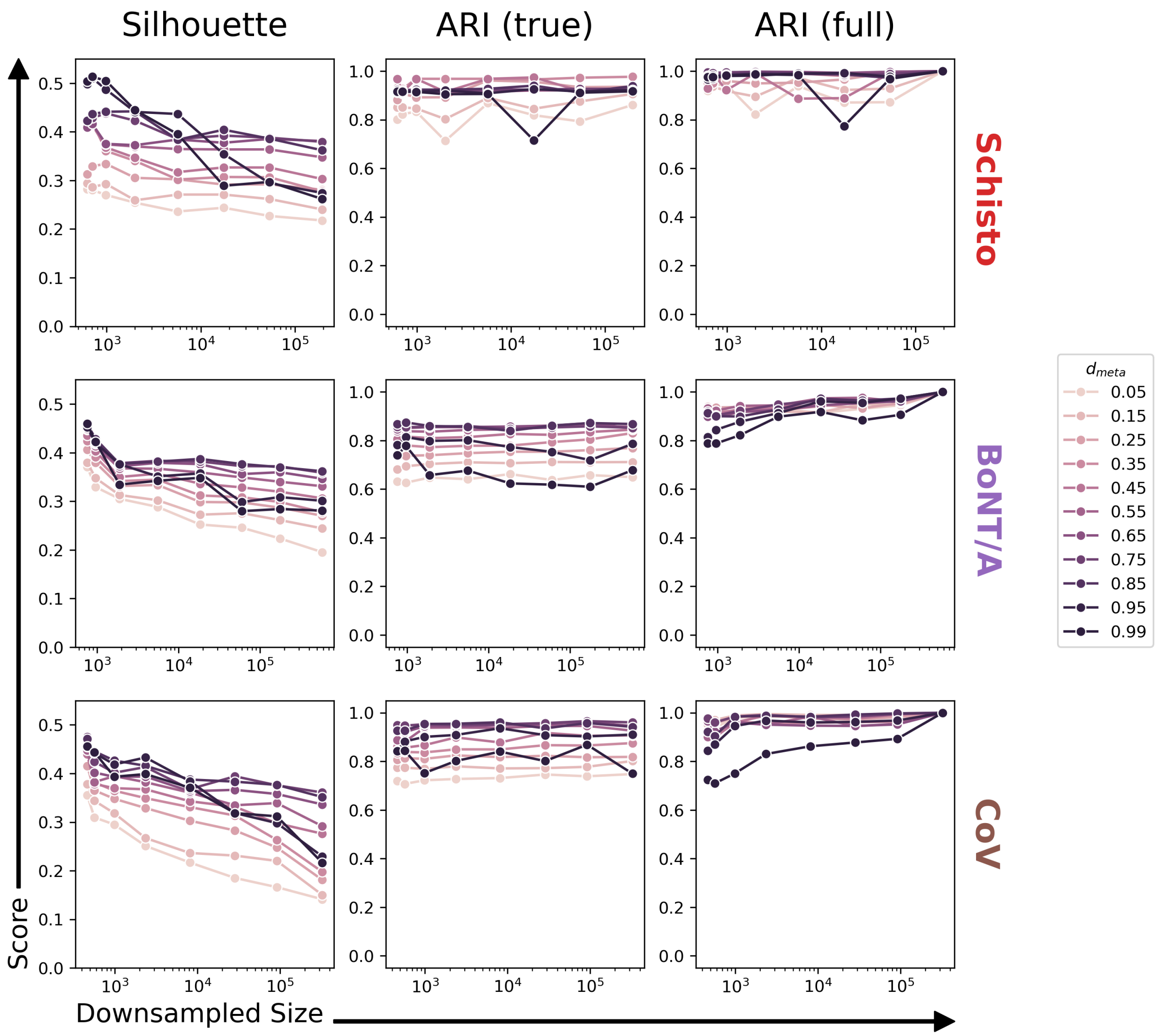


**Supplementary Figure 6: Sensitivity of average-linkage meta-clustering to dataset size.** Plots as in **figure S5** showing results for meta-clustering with $M_{meta}$ = average, and varying values of $d_{meta}$.

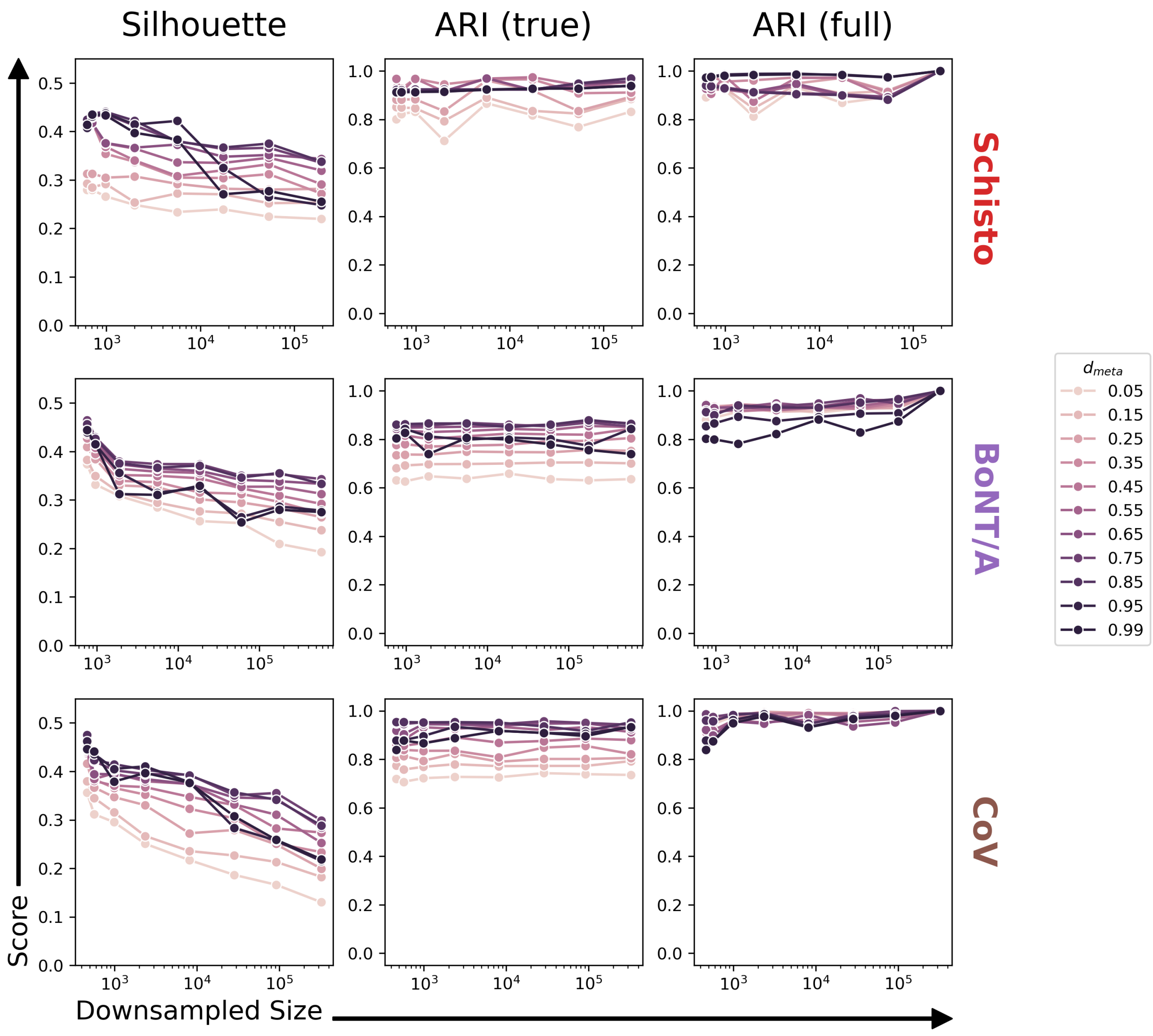


**Supplementary Figure 7: Sensitivity of complete-linkage meta-clustering to dataset size.** Plots as in **figure S5** showing results for meta-clustering with $M_{meta}$ = complete, and varying values of $d_{meta}$.

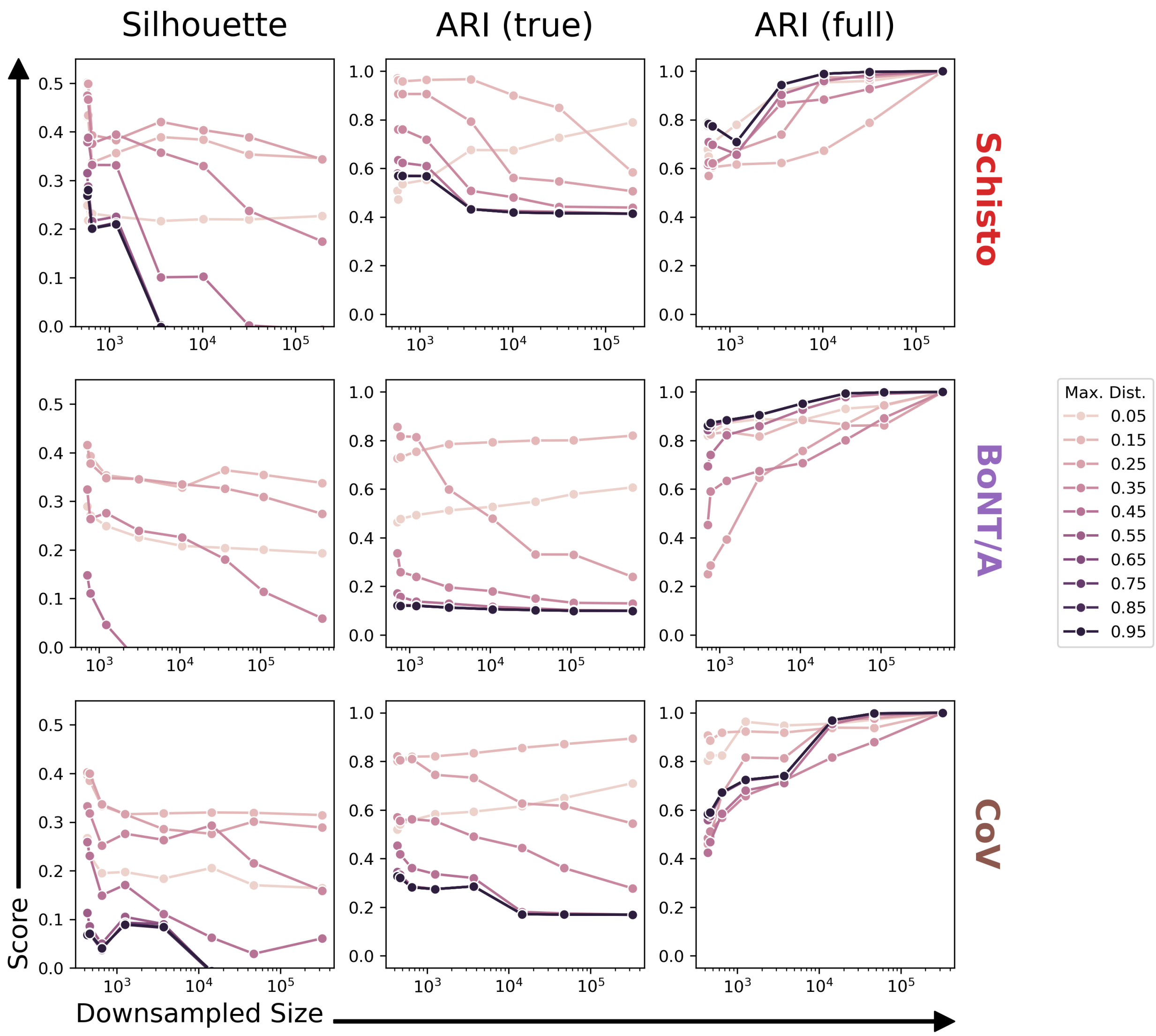


**Supplementary Figure 8: Sensitivity of single-linkage SCOPer clustering to dataset size.** Plots as in **figure S2** showing results for single-linkage SCOPer clustering, and varying distance cutoffs.

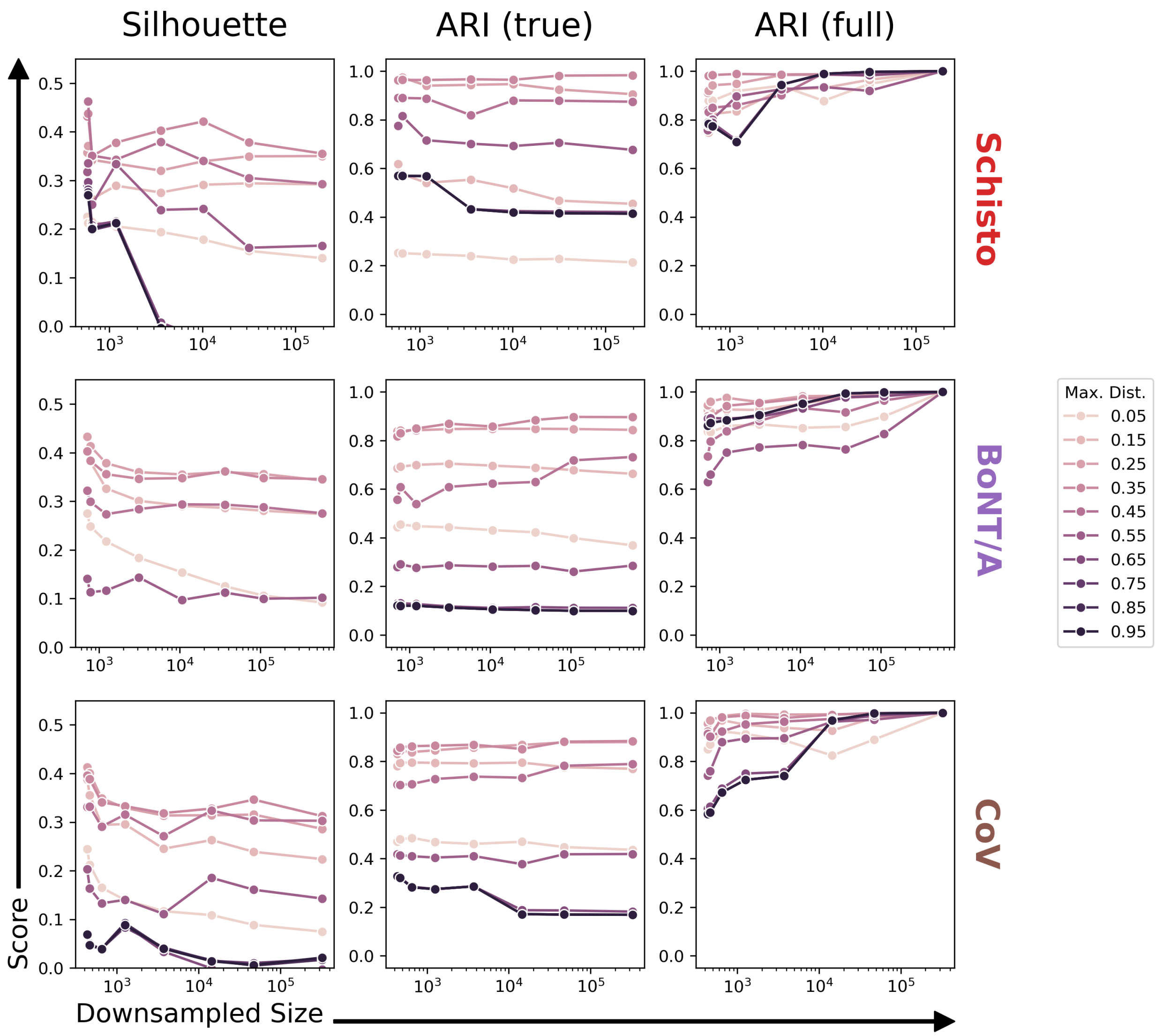


**Supplementary Figure 9: Sensitivity of average-linkage SCOPer clustering to dataset size.** Plots as in **figure S9** showing results for average-linkage SCOPer clustering.

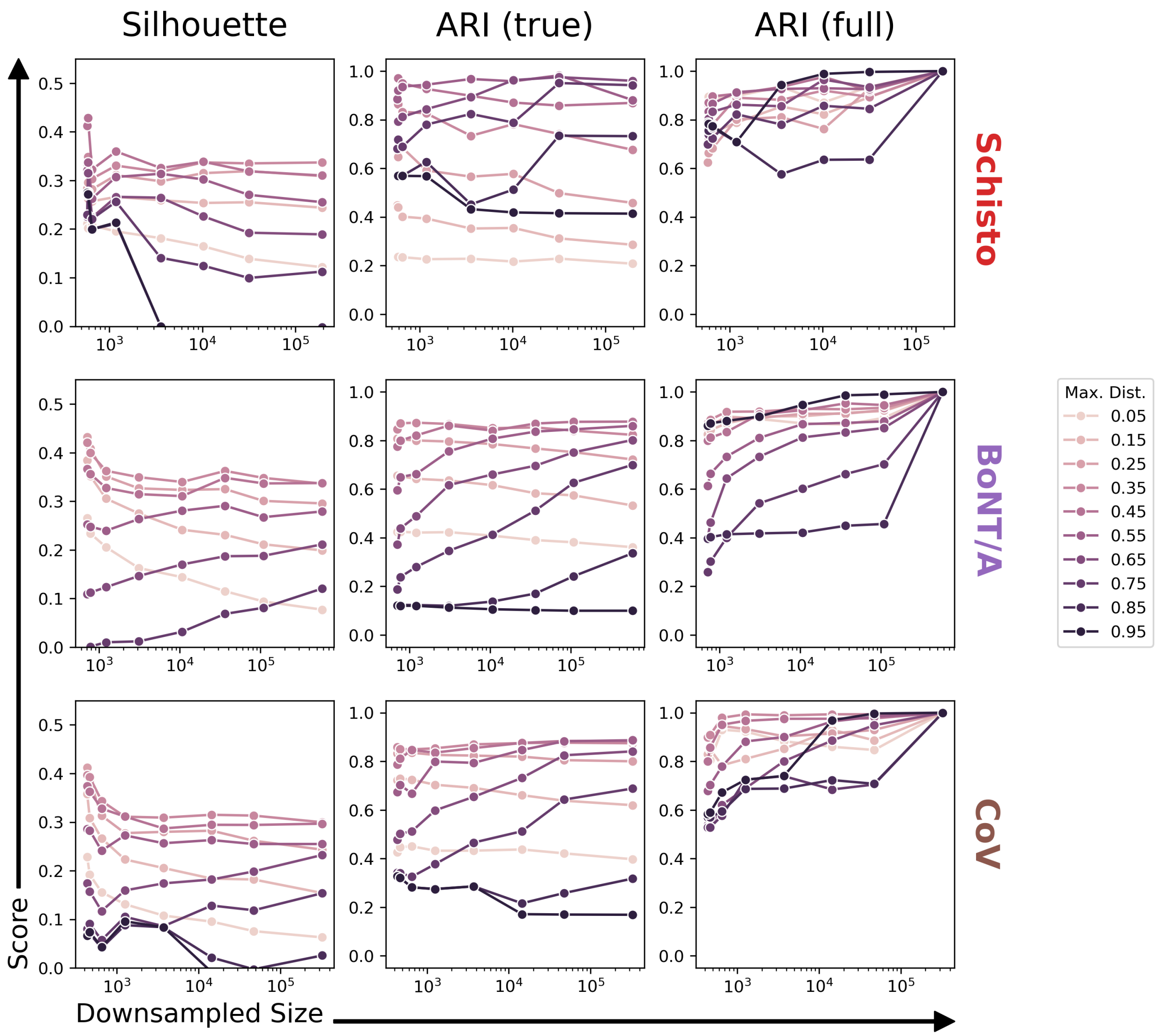


**Supplementary Figure 10: Sensitivity of complete-linkage SCOPer clustering to dataset size.** Plots as in **figure S9** showing results for complete-linkage SCOPer clustering.

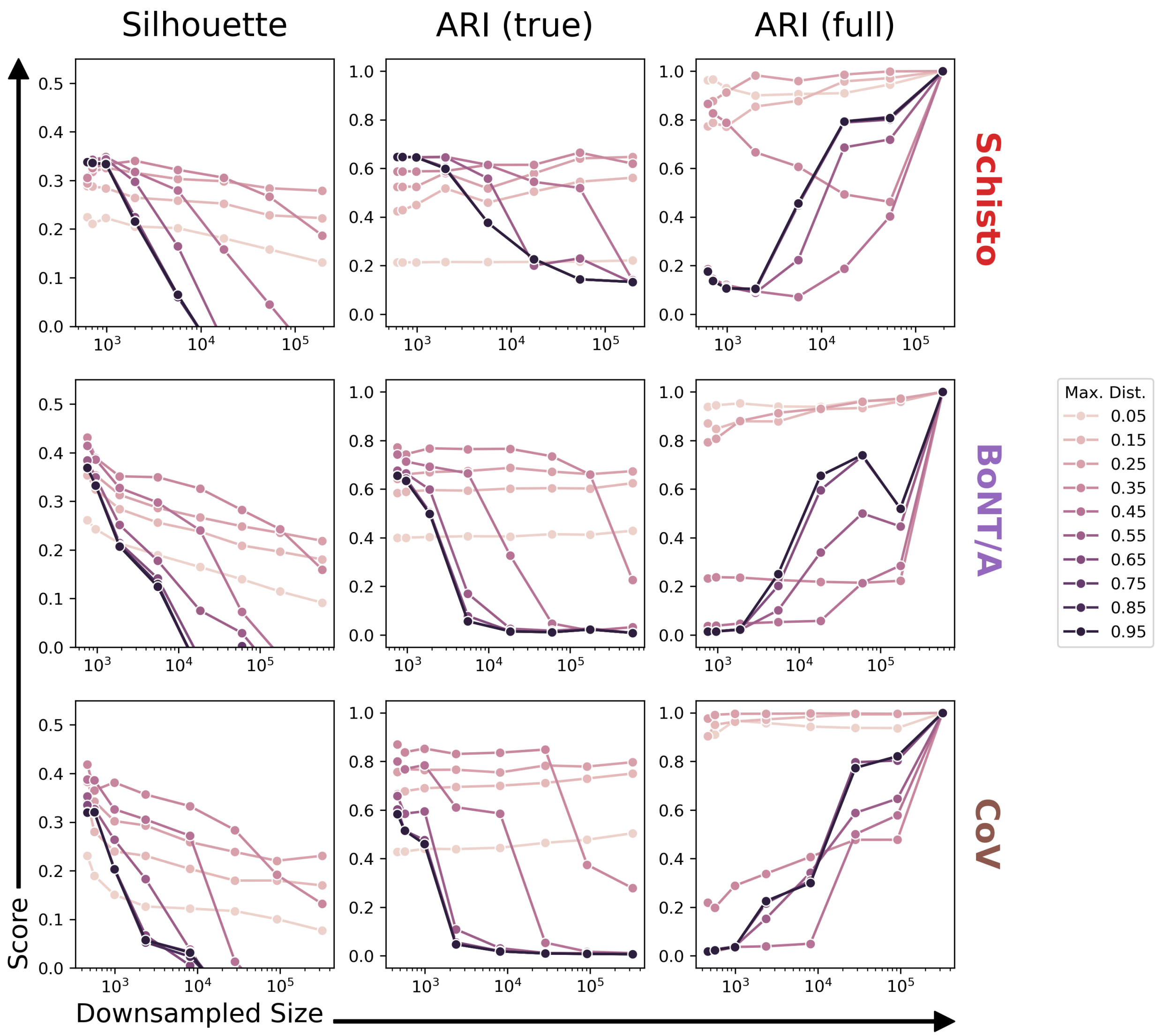


**Supplementary Figure 11: Sensitivity of connected component MMseqs2 clustering to dataset size.** Plots as in **figure S2** showing results for MMseqs2 clustering using the connected components clustering mode and varying distance cutoffs.

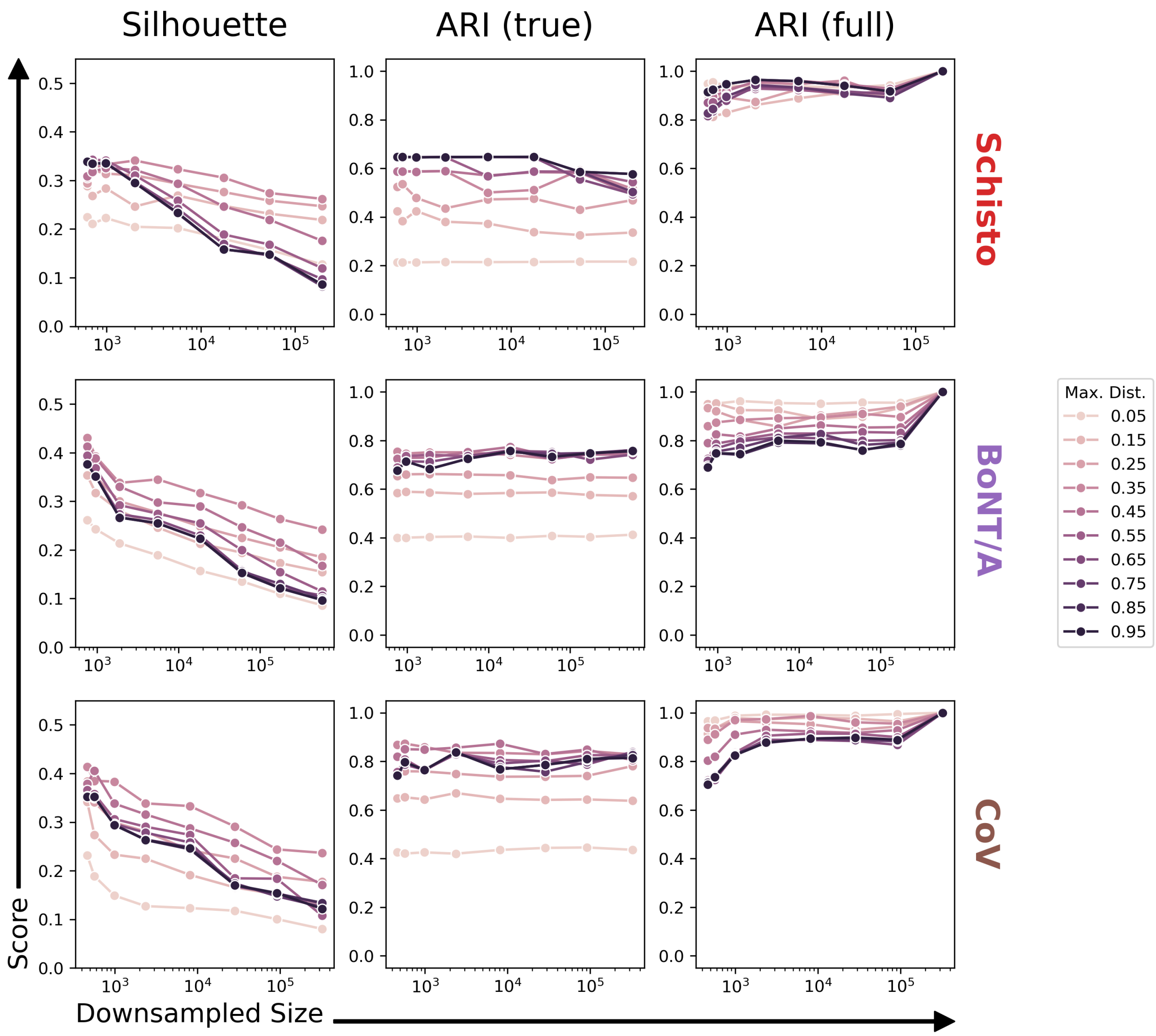


**Supplementary Figure 12: Sensitivity of greedy set cover MMseqs2 clustering to dataset size.** Plots as in **figure S11** showing results for MMseqs2 clustering using the greedy set cover clustering mode and varying distance cutoffs.

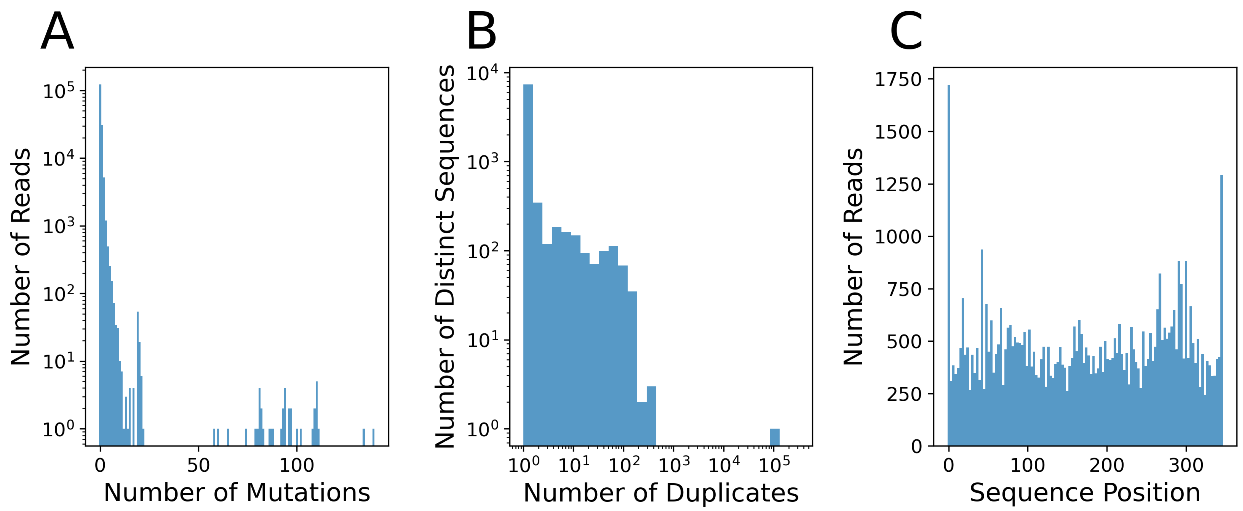


**Supplementary Figure 13: Error rates in DNA preparation.** Data from a sequencing experiment where a single nanobody sequence, derived from a clonal expression plasmid, was prepared and sequenced using the same PCR methods that were used for the cDNA nanobody repertoire sequencing. A) Histogram showing the distribution of the number of mutations over all sequencing reads. B) Histogram showing the distribution of duplicates over all distinct sequences observed. The bar with a height of 1 on the far right of the plot corresponds to an exact match to the sequence of the plasmid that was used in this experiment. C) Bar plot showing the number of reads containing a mutation for each position in the sequence. Deletions are counted as mutations in the position that was deleted and insertions are counted in the position immediately preceding the insertion.

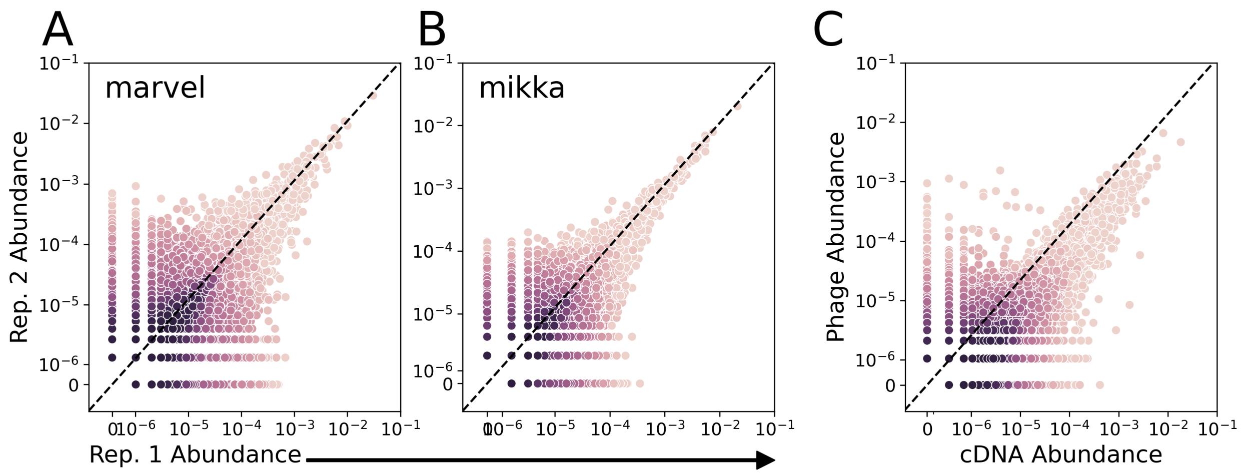


**Supplementary Figure 14: DNA library preparation and sequencing yield reproducible results.** Scatter plots comparing nanobody abundances (expressed as the fraction of total reads corresponding to a specific amino acid sequence) between various pairs of samples. Points are colored by the log of the density of nearby points, with darker points indicating higher density. Black dashed lines show the ideal relationship (a slope of 1, and intercept of 0). A,B) Comparing cDNA derived from two replicate blood draws taken from each of two alpacas, Marvel (A) and Mikka (B). C) Comparing cDNA abundance (averaged across both alpacas) to abundance in a nanobody display phage display library prepared from pooled cDNA of both alpacas.

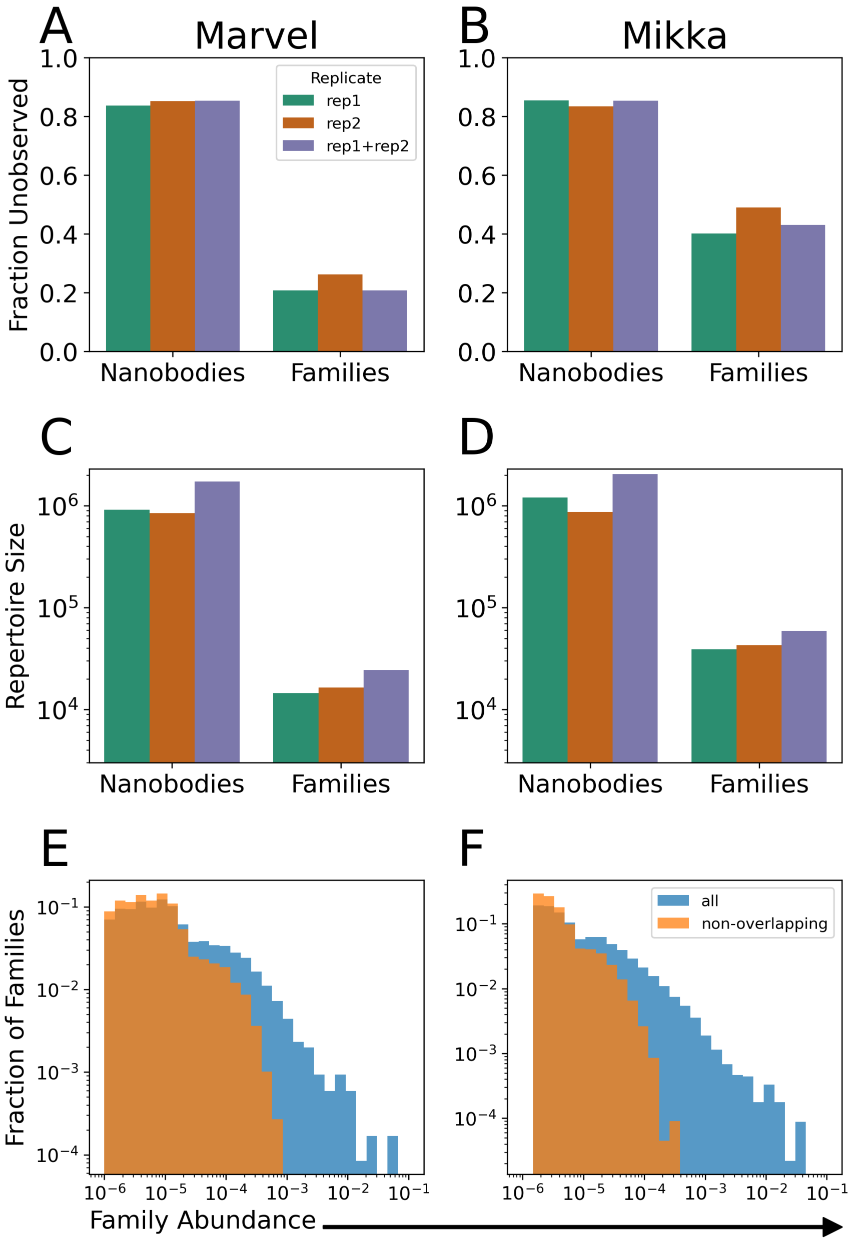


**Supplementary Figure 15: cDNA libraries capture a large portion of the alpaca immune repertoire.** Preseq estimations of repertoire size for the same replicate samples as in **figure S14**. Estimates are made separately for each replcate (rep1, rep2) or for pooled data across replicates (rep1+rep2). Repertore sizes are estimated at the amino acid sequence (nanobody) level, and at the family level (using NanoMAP individual clustering with the established optimal parameters). A,B) Estimates of the fraction of total families (or nanobodies) that were left unobserved in our sequencing experiments for either Marvel (A) or Mikka (B). C,D) Estimates of the total number of families (or nanobodies) in the repertoire of either Marvel (C) or Mikka (D). E,F) Abundance distributions for clonal families clustered with $d_{meta} = 0.75$ and $M_{meta} =$ single, showing the distribution for all families (blue) or only those families that appear in one replicate but not the other (orange). Family abundance is defined as the fraction of total reads mapped to the family.

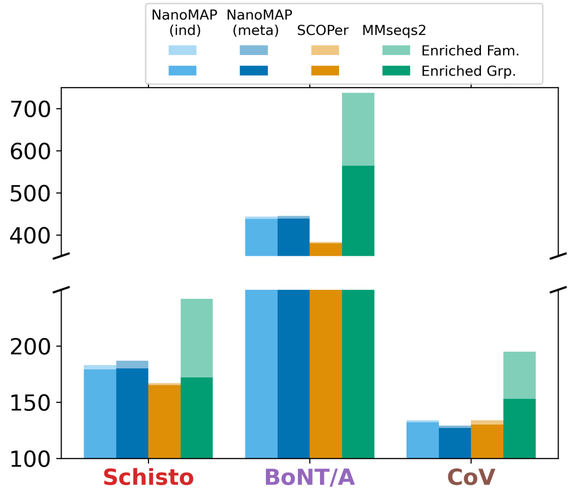


**Supplementary Figure 16: NanoMAP clusterings result in more distinct candidate families.** Bar chart showing the number of families in each dataset identified as candidates for binding to at least one target by each clustering method. Parameters for each method are the same as in **Fig. 3**. Families with GFold scores above 2 for a target are considered to be candidates. Light bars indicate the total number of candidate families, while light bars indicate the number of groups of related families. Groups of related families are identified as described in **section S1.11**.

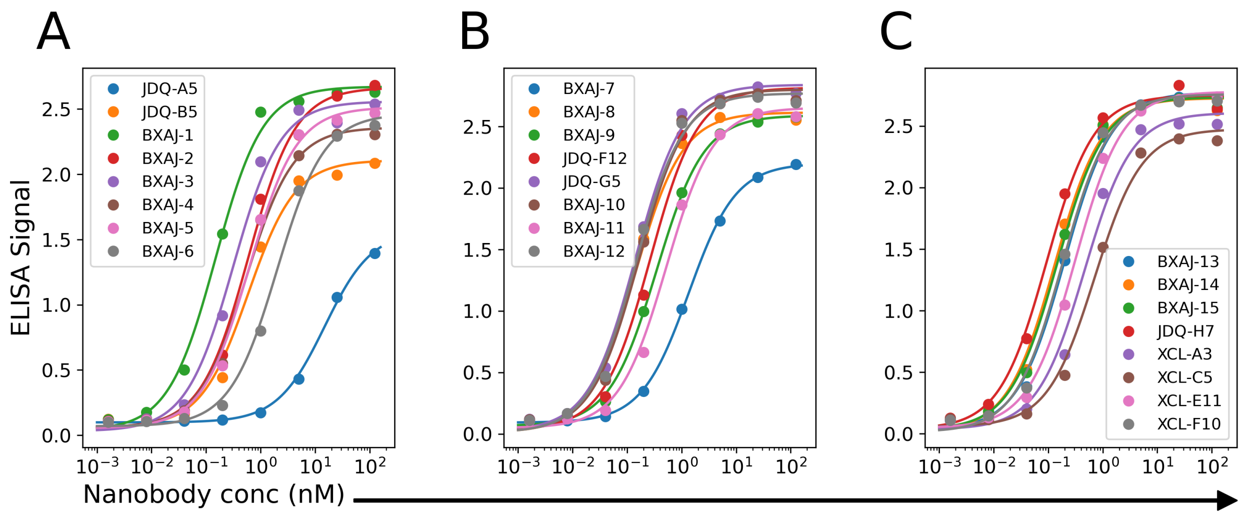


**Supplementary Figure 17: BoNT/A dilution ELISAs.** A-C) ELISA binding measurements (dots) and fitted binding curves (lines) for each of 24 selected nanobodies from the BoNT/A dataset. Each panel shows data for a different set of 8 nanobodies of the 24 named in the legends.

### References

[1] T. Daley and A. D. Smith. Predicting the molecular complexity of sequencing libraries. *Nature methods*, 10(4):325–327, 2013.

[2] P. Dash, A. J. Fiore-Gartland, T. Hertz, G. C. Wang, S. Sharma, A. Souquette, J. C. Crawford, E. B. Clemens, T. H. Nguyen, K. Kedzierska, et al. Quantifiable predictive features define epitope-specific T cell receptor repertoires. *Nature*, 547(7661):89–93, 2017.

[3] F. Ehrenmann and M.-P. Lefranc. IMGT/DomainGapAlign: IMGT standardized analysis of amino acid sequences of variable, constant, and groove domains (IG, TR, MH, IgSF, MhSF). *Cold Spring Harbor Protocols*, 2011(6):pdb–prot5636, 2011.

[4] J. Feng, C. A. Meyer, Q. Wang, J. S. Liu, X. Shirley Liu, and Y. Zhang. Gfold: a generalized fold change for ranking differentially expressed genes from rna-seq data. *Bioinformatics*, 28(21):2782–2788, 2012.

[5] N. T. Gupta, J. A. Vander Heiden, M. Uduman, D. Gadala-Maria, G. Yaari, and S. H. Kleinstein. Change-O: a toolkit for analyzing large-scale B cell immunoglobulin repertoire sequencing data. *Bioinformatics*, 31(20):3356–3358, 2015.

[6] J. J. Jaskiewicz, J. M. Tremblay, S. Tzipori, and C. B. Shoemaker. Identification and characterization of a new 34 kDa MORN motif-containing sporozoite surface-exposed protein, Cp-P34, unique to Cryptosporidium. *International journal for parasitology*, 51(9):761–775, 2021.

[7] F. Madeira, N. Madhusoodanan, J. Lee, A. Eusebi, A. Niewielska, A. R. Tivey, R. Lopez, and S. Butcher. The EMBL-EBI Job Dispatcher sequence analysis tools framework in 2024. *Nucleic acids research*, 52(W1):W521–W525, 2024.

[8] N. Nouri and S. H. Kleinstein. Somatic hypermutation analysis for improved identification of B cell clonal families from next-generation sequencing data. *PLoS computational biology*, 16(6):e1007977, 2020.

[9] P. J. Rousseeuw. Silhouettes: a graphical aid to the interpretation and validation of cluster analysis. *Journal of computational and applied mathematics*, 20:53–65, 1987.

[10] M. Steinegger and J. Söding. Mmseqs2 enables sensitive protein sequence searching for the analysis of massive data sets. *Nature biotechnology*, 35(11):1026–1028, 2017.

[11] V. A. Traag, L. Waltman, and N. J. Van Eck. From louvain to leiden: guaranteeing well-connected communities. *Scientific reports*, 9(1):5233, 2019.
